# SIRT1 remodels astrocyte metabolism and promotes viral replication during neurotropic orthoflavivirus infection

**DOI:** 10.64898/2026.07.23.740149

**Authors:** Juan P. Angel, Diego Alzate, Marissa Lindman, Colm Atkins, Brian P. Daniels

## Abstract

Orthoflaviviruses depend on host metabolic resources to replicate, but how distinct cell types in the central nervous system (CNS) alter their metabolism in response to infection remains incompletely understood. Here, we examined whether the NAD+-dependent deacetylase SIRT1 is engaged during Zika virus (ZIKV) and West Nile virus (WNV) infection and whether its activity influences infection outcomes. SIRT1 activity increased in astrocytes isolated from infected mouse brains and in infected primary human astrocytes, but not in neurons or CNS myeloid cells. In astrocytes, infection induced transcriptomic, metabolomic, and functional changes consistent with SIRT1 activation, enhanced NAD+ salvage, and enhanced oxidative metabolism. Pharmacologic inhibition of SIRT1 or NAD+ salvage reduced ZIKV replication in astrocytes, whereas SIRT1 activation or supplementation with an NAD+ precursor increased replication. In mice, SIRT1 inhibition or nicotinamide (NAM) treatment reduced viral burden and mortality following ZIKV and WNV infection, while SIRT1 activation worsened disease. Together, these findings identify NAD+/SIRT1 as key regulators of cellular metabolism in infected astrocytes and support a role for this pathway in promoting orthoflavivirus replication and pathogenesis.

## Introduction

Orthoflaviviruses, such as Zika virus (ZIKV) and West Nile virus (WNV), are neurotropic pathogens that can cause encephalitis and other neurologic complications. The recent expansion of several orthoflavivirus genera across the Americas and Europe underscores their growing threat to global public health (1–3). Within the central nervous system (CNS), disease outcome depends not only on viral tropism and host immunity, but also on the physiologic state of infected cells. Different CNS cell populations vary substantially in their permissiveness to infection, antiviral capacity, and ability to tolerate the metabolic demands imposed by viral replication (4–7).

Orthoflaviviruses rely on host metabolic pathways to generate the energy and biosynthetic intermediates required for replication. Infection can alter glycolysis, mitochondrial respiration, nucleotide synthesis, lipid metabolism, and redox balance, but the direction and consequence of this reprogramming are strongly cell-type dependent (8, 9). In fibroblasts, endothelial cells, macrophages, and other non-neural populations, ZIKV and WNV infection commonly promote aerobic glycolysis, and inhibition of glycolytic or nucleotide-generating pathways can restrict replication in these cell types (10–13). In contrast, we and others have shown that a shift into glycolysis is associated with an antiviral state in neurons of the CNS (14). Studies of isolated CNS cell populations suggest that CNS glial cells may exhibit metabolic responses distinct from those of neurons and non-CNS cell types, but the metabolic programs that support or restrict viral replication in these cells remain incompletely understood (15–17).

Astrocytes, the most abundant glial cell type in the CNS, are well positioned to coordinate metabolic and antiviral responses during CNS infection. They maintain a predominantly glycolytic metabolic baseline but retain functional mitochondria and substantial metabolic flexibility, allowing them to adjust between glycolysis and oxidative phosphorylation in response to cellular stress (18, 19). Astrocytes also mount strong innate immune responses to ZIKV and WNV, including type I interferon signaling and production of inflammatory chemokines (4, 20, 21). These metabolic and immune functions are closely linked, as inflammatory signaling can reshape mitochondrial and glycolytic activity, while metabolite availability influences the transcriptional programs that govern antiviral defense (22). Despite these insights, the mechanisms that regulate metabolic state in orthoflavivirus-infected astrocytes, and the consequences of that regulation for viral replication, remain poorly defined.

Sirtuin 1 (SIRT1) is a candidate regulator of this response due to its overlapping regulatory control of both cellular metabolism and transcription. SIRT1 is an NAD+-dependent deacetylase that modifies transcription factors involved in oxidative stress, inflammation, mitochondrial function, and energy metabolism, including FOXO proteins, p53, NF-κB, and HIF1A (23–26).

Depending on cellular context, SIRT1 can promote oxidative metabolism, regulate stress resistance, and alter innate immune signaling. It has also been implicated in control of IRF- and JAK-STAT-dependent antiviral responses, although its effects vary across experimental systems (27, 28). Thus, SIRT1 is positioned to influence both the metabolic environment available to replicating viruses as well as the host transcriptional response to infection.

However, whether SIRT1 is activated during orthoflavivirus infection in the CNS, and whether this response is cell-type specific, has not been well established.

Because SIRT1 consumes NAD+ during deacetylation, its activity depends on continued NAD+ regeneration. In mammalian cells, this is maintained primarily through the salvage pathway, in which nicotinamide phosphoribosyltransferase (NAMPT) converts nicotinamide (NAM) to nicotinamide mononucleotide (NMN), which is then converted to NAD+ by nicotinamide mononucleotide adenylyltransferase (NMNAT) enzymes (29–31). This pathway not only replenishes the SIRT1 cosubstrate but also clears NAM, a product of NAD+-dependent deacetylation that can inhibit sirtuin activity. Manipulation of NAMPT, NMN, or related salvage-pathway components can alter NAD+ abundance and SIRT1-dependent functions in astrocytes (32). Orthoflavivirus infection has also been associated with changes in NAD+-associated pathways, oxidative phosphorylation, and central carbon metabolism in the brain, raising the possibility that the NAD+/SIRT1 axis may contribute to infection-induced metabolic remodeling (33).

Here, we tested whether orthoflavivirus infection engages NAD+ salvage and SIRT1 activity in a cell-type-specific manner and whether this response influences viral replication and pathogenesis in models of adult orthoflavivirus encephalitis. We show that SIRT1 activity is selectively increased in infected astrocytes and is accompanied by transcriptional and metabolic changes consistent with enhanced NAD+ salvage and increased oxidative metabolism.

Pharmacologic manipulation of SIRT1 and the NAD+ salvage pathway produced reciprocal effects on viral replication in astrocytes and on disease severity in mouse models of ZIKV and WNV infection which, together, suggest that SIRT1 activity supports orthoflavivirus replication and disease pathogenesis. These findings identify the NAD+/SIRT1 axis as an important regulator of astrocytic metabolism and host defense during neurotropic orthoflavivirus infection.

## Results

### Orthoflavivirus infection induces SIRT1 activation in astrocytes but not in neurons or CNS myeloid cells

Sirtuins exert cell-type-specific functions, and astrocytes, neurons, and CNS myeloid cells differ in baseline metabolic state, NAD+ availability, and susceptibility to infection. We thus asked whether SIRT1 activity is differentially induced across these populations during orthoflavivirus infection. We used two complementary models of orthoflavivirus encephalitis to address this question. In the first, wildtype C57BL/6J mice were intracranially inoculated with ZIKV MR766 (10^4^ PFU) and brains were harvested at 4 days post-infection. In the second, C57BL/6J mice were inoculated in the footpad with WNV-Bird114 (10^3^ PFU) and brains were harvested at 9 days post-infection. In both models, brains were subjected to enzymatic dissociation and magnetic-activated cell sorting (MACS) to isolate distinct CNS cell populations (Fig. 1A).

**Figure 1.**
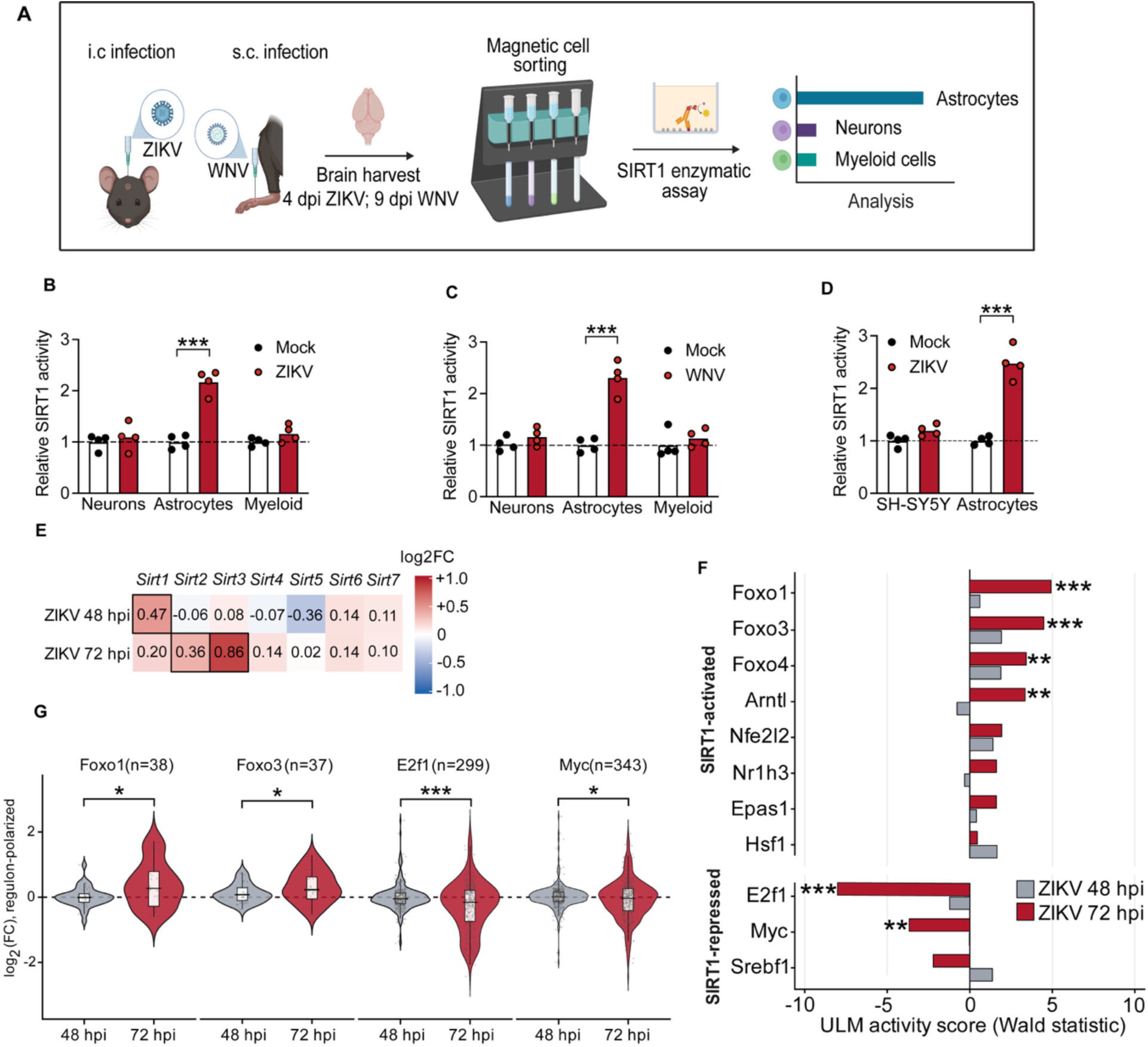
Flavivirus infection induces SIRT1 activation in astrocytes but not in neurons or CNS myeloid cells. (A) Schematic of the two encephalitis models used in this study. C57BL/6J mice were inoculated intracranially with ZIKV MR766 (10⁴ PFU) with brains harvested at 4 dpi, or in the footpad with WNV-Bird114 (10³ PFU) with brains harvested at 9 dpi, followed by magnetic-activated cell sorting (MACS) of indicated cell populations. This panel was created with BioRender.com. (B, C) SIRT1 deacetylase activity in indicated cell types sorted from brains of mice infected with ZIKV (B) or WNV (C). n = 4 mice/group. Comparisons via Welch’s two-tailed t-test. (D) SIRT1 deacetylase activity in ZIKV-infected primary human astrocytes and differentiated SH-SY5Y cultures at 48 hpi. n = 3 independent cultures/group. Comparisons via Welch’s two-tailed t-test. (E) Log₂ fold change in sirtuin-encoding gene expression in ZIKV-infected primary mouse astrocytes at 48 and 72 hpi relative to mock controls. n = 3 cultures/timepoint. Cells outlined in black indicate padj < 0.05. (F) Transcription factor activity for the 11 SIRT1-substrate regulons, inferred by decoupleR (univariate linear model) from mouse DoRothEA regulons, comparing ZIKV-infected astrocytes to mock controls. Comparisons via Benjamini-Hochberg procedure. (G) Per-gene log₂ fold change distributions for indicated regulons at 72 hpi, shown as violin plots with overlaid box plots (median and interquartile range). Comparisons via paired Wilcoxon signed-rank test, with pre-ranked gene set enrichment analysis (GSEA) against the 72 hpi Wald statistic across all 11 substrate regulons (with Benjamini-Hochberg correction). \**p* < 0.05, \*\**p* < 0.01, \*\*\**p* < 0.001.

We next assayed SIRT1 deacetylase activity in astrocytes, neurons, and myeloid cells sorted from mock- and orthoflavivirus-infected brains using a fluorometric assay in which SIRT1-dependent deacetylation of a fluorogenic substrate produces a measurable fluorescence signal corresponding to the magnitude of active SIRT1 in each sample. We observed that SIRT1 activity was significantly elevated in sorted astrocytes in both infection models (Fig. 1B-C). In contrast, neurons and myeloid cells showed no detectable change in SIRT1 activity following infection with either virus. To determine whether SIRT1 activation also occurs in orthoflavivirus-infected human cells, we infected “neuron-like” differentiated SH-SY5Y neuroblastoma cells and primary human astrocytes with ZIKV MR766 (MOI 0.01) and measured SIRT1 deacetylase activity using the same fluorometric assay at 48 hours post-infection (hpi). ZIKV infection increased SIRT1 activity in primary human astrocytes, but not in differentiated SH-SY5Y cultures (Fig. 1D). Together, these findings suggest that astrocytes are the principal CNS cell population in which SIRT1 activity is engaged during orthoflavivirus infection.

Having shown that orthoflavivirus infection elevates SIRT1 activity in astrocytes, we asked whether infection also drives *Sirt1* expression. We thus performed bulk RNA sequencing (RNAseq) in primary mouse astrocytes infected with ZIKV-MR766 (MOI 0.01), comparing expression of Sirtuin-encoding genes at 48 and 72 hpi relative to mock-infected controls. *Sirt1* was modestly upregulated at 48 hpi, while *Sirt2* and *Sirt3* were elevated at 72 hpi (Fig. 1E). Transcript levels of other Sirtuin enzymes, including those encoded by *Sirt4, Sirt5, Sirt6,* and *Sirt7*, were not statistically distinguishable from control levels at either time point. These data suggest that ZIKV infection results in both increased transcript expression and increased enzymatic activity of SIRT1 in astrocytes, although the modest increase observed in transcript expression suggests that other mechanisms may underly the observed enhancement in SIRT1 enzymatic activity.

Given the important roles for SIRT1 in regulating cellular transcription, we next asked whether ZIKV infection produced gene-expression changes consistent with increased SIRT1 activity in astrocytes by using decoupleR to estimate the activities of transcription factors regulated by SIRT1. This approach assesses whether the established target genes, or regulon, of each transcription factor show coordinated changes in expression. We compared ZIKV-infected astrocytes with mock-infected controls at 48 and 72 hpi using mouse DoRothEA regulons for 273 transcription factors. Notably, SIRT1 is reported to enhance the activity of FOXO1, FOXO3, FOXO4, ARNTL, NFE2L2, NR1H3, EPAS1, and HSF1, while inhibiting or destabilizing E2F1, MYC, and SREBF1 (34, 35). We therefore interpreted increased activity of the first group and decreased activity of the second as a transcriptional signature consistent with SIRT1 activation. At 72 hpi, the inferred activities of all 11 factors changed in the predicted direction: the eight positively regulated factors increased, whereas the three negatively regulated factors decreased. Six of the 11 factors reached statistical significance, with FOXO1, FOXO3, FOXO4, and ARNTL activity increased and E2F1 and MYC activity decreased (Fig. 1F). By contrast, none of the 11 factors reached significance at 48 hpi. Thus, ZIKV infection induces a coordinated transcriptional signature consistent with increased SIRT1 activity between 48 and 72 hpi.

To assess whether these inferred activity scores were supported by coordinated changes among the underlying target genes, rather than being specific to the decoupleR scoring method, we also examined gene-level expression changes within the four regulons large enough to support analysis of per-gene distributions: FOXO1, FOXO3, E2F1, and MYC (Fig. 1G). At 48 hpi, the distributions of fold change expression values for target genes in each regulon were centered near zero, indicating little coordinated regulation. However, by 72 hpi, each regulon shifted in the direction predicted by increased SIRT1 activity, with FOXO1 and FOXO3 target genes preferentially upregulated and E2F1 and MYC target genes preferentially downregulated. Paired Wilcoxon tests comparing fold change values for the same target genes at 48 and 72 hpi identified significant shifts for all four regulons, with the largest effect in E2F1. As an additional analysis using a distinct enrichment method, we performed pre-ranked gene set enrichment analysis (GSEA) against the 72 hpi Wald statistic across all 11 substrate regulons (Fig. S1A). This analysis identified significant negative enrichment of the E2F1 and MYC regulons. The remaining substrate regulons showed enrichment in the predicted direction, with FOXO4 and ARNTL reaching statistical significance. Together, these analyses indicate that the inferred activity changes in SIRT1-associated regulons are reflected in coordinated expression patterns of the corresponding target genes and are not an artifact of a single scoring approach.

As an additional analysis using a pathway database independent of DoRothEA, we analyzed the same RNA-seq dataset with Ingenuity Pathway Analysis (IPA). IPA identified enrichment of several pathways associated with established SIRT1-regulated processes, including FOXO transcriptional signaling (23, 36, 37), NRF2 antioxidant signaling (38, 39), FOXO-mediated microautophagy (40), PGC1α-associated mitochondrial division (41), as well as NAD signaling and nicotinate metabolism (Fig. S1B). In contrast, the broader Sirtuin Signaling pathway showed little net predicted activation or inhibition, likely reflecting the inclusion of multiple sirtuin family members and downstream processes with opposing directional changes. Together with our regulon-based analyses, these results indicate that ZIKV infection engages specific transcriptional programs associated with SIRT1 activity rather than producing uniform activation of the entire sirtuin signaling network. This coordinated pattern was most evident at 72 hpi, when the target signatures of SIRT1-activated factors were induced and those of SIRT1-repressed factors were suppressed.

To determine whether this SIRT1-associated transcriptional program was specific to astrocytes among CNS cell types, we performed secondary analysis of a transcriptomic dataset previously generated by our group and others using primary mouse cortical neurons infected with ZIKV- MR766 or WNV-TX 2002-HC (14). We note that we could not precisely match the time interval post infection in our transcriptomic analysis of neurons and astrocytes because orthoflavivirus infection progresses differently in these cell types. Viral replication proceeds more slowly in astrocytes, which also remain viable for longer periods after infection, allowing transcriptional responses to be examined at 48 and 72 hpi. In contrast, orthoflaviviruses replicate more rapidly and induce substantial apoptosis in neurons (42, 43), making analysis at later time points increasingly confounded by selective cell loss. We therefore analyzed neurons at 24 hpi, which we considered a conceptually comparable stage of established infection before widespread neuronal death, rather than a strictly chronologically matched time point. Using the same regulon-based analysis pipeline, no SIRT1-regulated substrate transcription factor reached statistical significance in neurons infected with either virus (Fig. S1C). Consistent with this result, IPA Upstream Regulator analysis predicted activation of the SIRT1-regulated factors NRF2, PPARG, ESRRA, FOXO1, and ATG7 in infected astrocytes, while predicting inhibition of these factors in infected neurons (Fig. S1D). We also did not observe significant alteration to transcript abundance of any Sirtuin-encoding genes in neurons infected with either virus (Fig. S1E). These findings indicate that the coordinated SIRT1-associated transcriptional program observed in astrocytes was not similarly engaged in neurons during the viable phase of established ZIKV or WNV infection. Nevertheless, because our neuronal analysis was limited to 24 hpi, we cannot exclude the possibility that a comparable signature emerges later in the subset of neurons that survives infection. Together, these findings suggest that orthoflavivirus infection selectively increases SIRT1 activity in astrocytes, but not neurons or CNS myeloid cells. In astrocytes, this response is accompanied by a delayed, coordinated transcriptional program consistent with SIRT1 activation, including activation of FOXO-associated pathways and repression of E2F1- and MYC-associated programs, suggesting significant functional implications for cellular metabolism and host defense.

### ZIKV infection remodels astrocyte metabolism in a manner favorable to sustained SIRT1 activity by preserving NAD+ and limiting accumulation of NAM

We previously established that orthoflavivirus infection increases SIRT1 deacetylase activity in astrocytes (Fig. 1B-D). Because SIRT1 requires NAD+ as a cosubstrate, SIRT1 activity is influenced by the availability and turnover of the cellular NAD pool. During each deacetylation reaction, SIRT1 consumes NAD+, which must be replenished to sustain continued activity. In astrocytes, NAD+ can be regenerated through the salvage pathway, in which NAM is converted to NMN by NAMPT. NMN is subsequently converted to NAD+ by NMNAT enzymes (Fig. 2A). We therefore asked whether ZIKV-MR766 infection altered astrocyte NAD metabolism by performing targeted metabolomics in primary human astrocytes at 48 hpi. NAD+ abundance was unchanged between mock- and ZIKV-infected cells (Fig. 2B). NADH and the total NAD pool showed non-significant downward trends (Fig. 2C-D), while the NAD+/NADH ratio showed a non-significant upward trend (Fig. 2E). Thus, despite increased SIRT1 activity, infection did not measurably reduce the cellular NAD+ pool at this time point.

**Figure 2.**
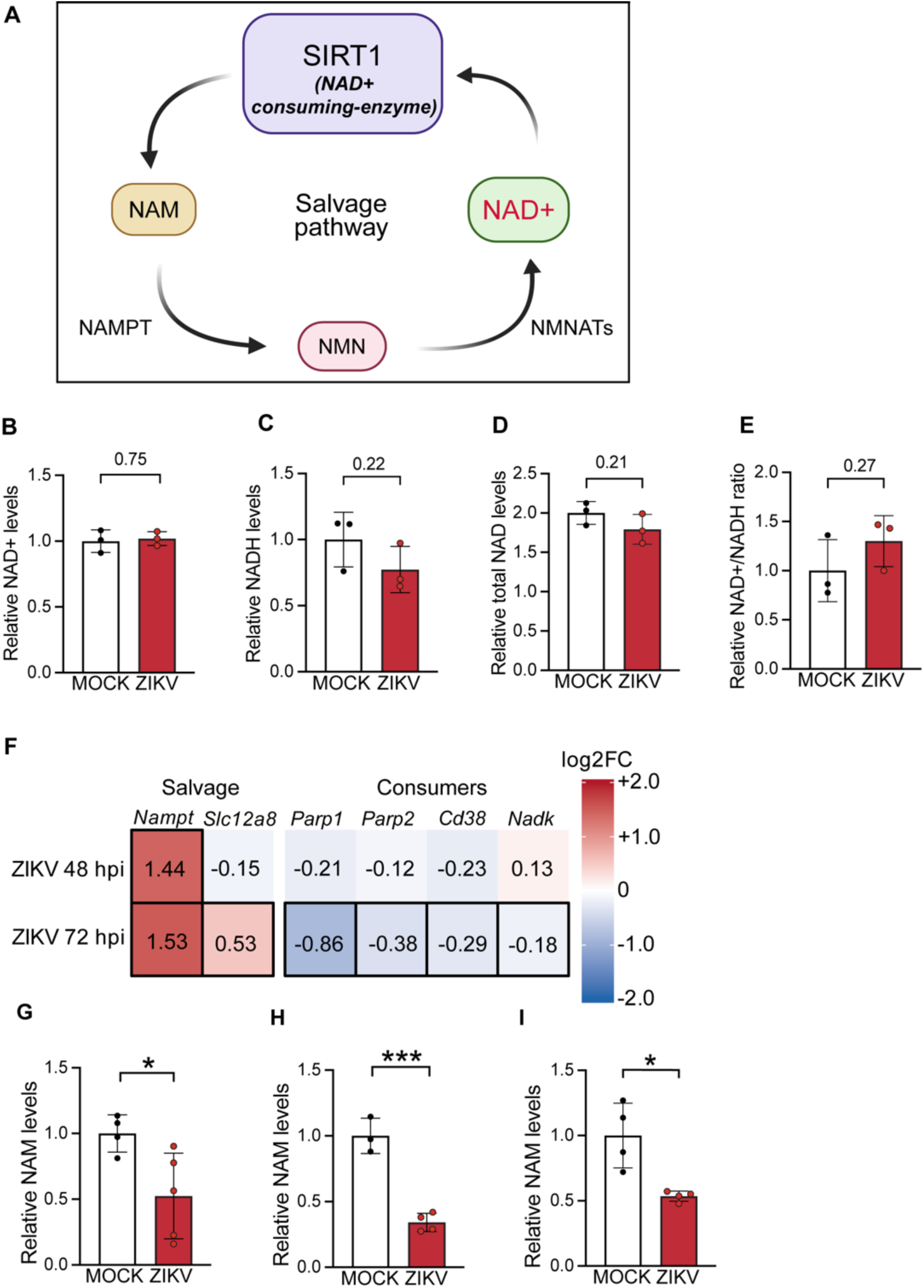
ZIKV infection remodels astrocyte metabolism in a manner favorable to sustained SIRT1 activity by preserving NAD+ and limiting accumulation of NAM. (A) Schematic of the NAD+ salvage pathway. This panel was created with BioRender.com. (B–E) Abundance of indicated metabolites in ZIKV-infected primary human astrocytes at 48 hpi, measured via LC-MS. Total NAD in (D) is the sum of individually normalized NAD+ and NADH. n = 3 independent cultures/group. Comparisons via Welch’s two-tailed t-test. (F) Log₂ fold change in expression of indicated genes in ZIKV-infected primary mouse astrocytes at 48 and 72 hpi relative to mock, (n = 3 cultures/ timepoint). Cells outlined in black indicate padj < 0.05. (G-I) NAM abundance in whole-brain homogenates from ZIKV-infected mice at 4 dpi (G), MACS-sorted astrocytes from ZIKV-infected mouse brains at 4 dpi (H), and ZIKV-infected primary human astrocytes at 72 hpi (I), measured by ELISA. For (G–I), NAM concentrations were normalized to total protein in each sample. Comparisons via Welch’s two-tailed t-test. Data in (B-E, and G-I) represent mean values ± SEM. \**p* < 0.05, \*\*\**p* < 0.001.

Our observation of a stable NAD+ pool during infection prompted us to examine transcriptional regulation of pathways that generate, consume, or redirect NAD+. We analyzed expression of genes in the NAD+ metabolic network by RNA-seq in primary mouse astrocytes at 48 and 72 hpi (Fig. 2F). The salvage-associated genes *Nampt* and *Slc12a8* were induced by infection, whereas several major NAD+-consuming enzymes, including *Parp1*, *Parp2*, and *Cd38*, were downregulated. *Nadk*, which diverts NAD+ toward NADP+ synthesis, was also modestly reduced. *Nampt*, which encodes the rate-limiting enzyme of the canonical salvage pathway, showed the strongest induction. This transcriptional remodeling developed over time, as *Nampt* was the only gene significantly altered at 48 hpi, while a broader metabolic program became apparent by 72 hpi, in parallel to the delayed emergence of the SIRT1-associated transcriptional program described in Fig. 1. The same coordinated pattern was not observed in primary mouse neurons analyzed at 24 hpi (Fig. S2A). Although *Cd38* was induced following ZIKV infection and *Nampt* was induced following WNV infection, neurons did not show the combined increase in salvage-associated genes and decrease in competing NAD+-consuming pathways observed in astrocytes. These data are therefore consistent with a model in which ZIKV infection remodels astrocyte NAD+ metabolism in a direction that may help preserve NAD+ availability for SIRT1 and other NAD+-dependent processes. In contrast, this response was not observed in neurons under the conditions examined.

Because NAMPT catalyzes the conversion of NAM to NMN, its induction suggested that infection might increase NAM utilization in astrocytes through the salvage pathway. Consistent with this possibility, NAM abundance was reduced in whole-brain homogenates 4 days after intracranial ZIKV infection (Fig. 2G) and in astrocytes sorted from infected brains at the same time point (Fig. 2H), localizing this change at least in part to the astrocyte compartment. NAM was likewise reduced in ZIKV-infected primary human astrocytes at 72 hpi (Fig. 2I). A similar reduction in NAM abundance was detected in whole-brain homogenates 9 days following peripheral WNV infection and in cultured mouse astrocytes infected with either ZIKV or WNV (Fig. S2B-E), suggesting that this response was not restricted to infection with a single orthoflavivirus. In contrast, NAM showed a non-significant trend toward increased abundance in ZIKV-infected primary mouse cortical neurons at 24 hpi (p = 0.06), consistent with the absence of a coordinated salvage-pathway response in these cells (Fig. S2F). We note that NAM is generated by NAD+-consuming reactions, including SIRT1-mediated deacetylation, and can inhibit SIRT1 through product feedback. Its reduction in infected astrocytes is therefore consistent with increased clearance through the salvage pathway, although metabolite abundance alone does not directly measure pathway flux. Considered together, the maintenance of NAD+ levels (Fig. 2B-E), induction of salvage-associated genes (Fig. 2F), and reduction in NAM (Fig. 2G-I) support a model in which infection remodels astrocyte NAD metabolism in a manner favorable to sustained SIRT1 activity, preserving its NAD+ cosubstrate while limiting accumulation of an inhibitory reaction product.

### ZIKV infection in astrocytes results in transcriptomic and metabolomic signatures consistent with enhanced oxidative phosphorylation

Because SIRT1 can promote mitochondrial biogenesis and oxidative metabolism while restraining glycolytic reprogramming, we next asked whether its increased activity in infected astrocytes was accompanied by transcriptional changes consistent with a shift toward oxidative phosphorylation. RNA-seq analysis in ZIKV-infected primary astrocytes revealed broad induction of genes associated with oxidative phosphorylation by 72 hpi, including genes spanning all five respiratory-chain complexes (Fig. 3A). Genes encoding multiple subunits of Complex I exhibited widespread upregulation, while some of the largest increases occurred among Complex IV component-encoding genes, including *Cox4i2* and *Cox7a1*. Several glycolytic genes were also induced, including the glucose transporter *Slc2a1*, *Hk2*, aldolase-family genes, and *Eno2* (Fig. 3B). In contrast, *Ldha* and *Pdk1*, which promote pyruvate conversion to lactate and restrict its entry into mitochondrial oxidation, remained near mock-infected levels. Thus, the astrocytic transcriptional response to ZIKV infection featured a combination of increased expression of respiratory machinery and genes associated with proximal glycolytic capacity but did not show the coordinated induction of lactate-producing genes typically associated with aerobic glycolysis. TCA-cycle associated genes were generally reduced (Fig. 3C). In contrast, primary mouse cortical neurons exhibited a notably distinct transcriptional response to ZIKV infection (Fig. S3A-C). *Ldha* and *Pdk1*, which remained unchanged in astrocytes, were significantly induced in infected neurons, whereas the broad increase in genes associated with oxidative phosphorylation observed in astrocytes was not detected. These data therefore suggest distinct, cell-type-specific metabolic programs during infection, in which astrocytes develop a transcriptional signature favoring oxidative capacity, whereas neurons develop a signature more consistent with diversion of pyruvate toward lactate.

**Figure 3.**
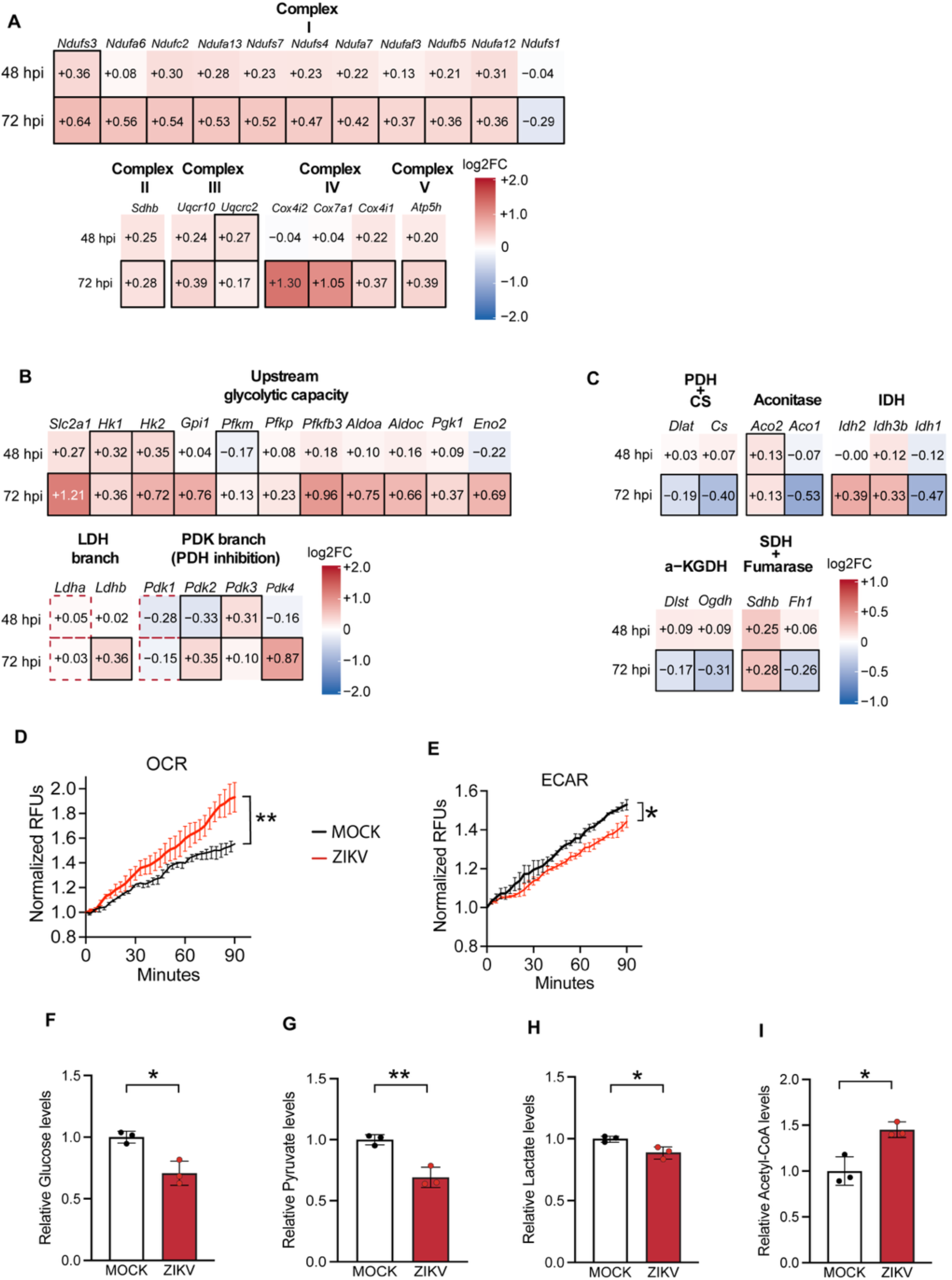
ZIKV infection in astrocytes results in transcriptomic and metabolomic signatures consistent with enhanced oxidative phosphorylation. (A-C) Log₂ fold change in expression of indicated genes in ZIKV-infected primary mouse astrocytes at 48 and 72 hpi relative to mock controls (n = 3 cultures/ timepoint). Cells outlined in black indicate padj < 0.05. (D, E) Oxygen consumption rate (D) and extracellular acidification rate (E) in ZIKV-infected primary mouse cortical astrocytes at 48 hpi. Measurements began at 48 hpi and proceeded for 90 minutes to capture the indicated readout. n = 3 independent cultures/group. Comparisons via Welch’s two-tailed t-test. (F–I) Relative abundance of indicated metabolites in ZIKV-infected primary human astrocytes at 48 hpi, measured by LC-MS. Comparisons via Welch’s two-tailed t-test. Data in (D-I) represent mean values ± SEM. \**p* < 0.05, \*\**p* < 0.01.

We next asked whether the transcriptional shift toward oxidative metabolism was accompanied by corresponding functional phenotypes. In primary mouse astrocytes, ZIKV infection increased oxygen consumption and reduced extracellular acidification at 72 hpi relative to mock infected controls (Fig. 3D-E), consistent with increased oxidative phosphorylation and reduced glycolytic output. Metabolomic analysis in primary human astrocytes at 48 hpi broadly supported the same interpretation: glucose, pyruvate, and lactate were reduced, whereas acetyl-CoA was increased (Fig. 3 F-I). These changes are consistent with increased routing of glucose-derived carbon into mitochondrial oxidation rather than conversion to lactate. Aspartate and glutamate, which are derived from the TCA cycle intermediates oxaloacetate and α-ketoglutarate and contribute to protein and nucleotide biosynthesis, were also significantly increased (Fig. S3D-E). Together, these functional and metabolomic data are concordant with our interpretation of the transcriptional program identified in Fig. 3A-C.

Primary mouse cortical neurons showed a contrasting metabolic response at 24 hpi. Infection induced *Pdk1* and *Pdk4* (Fig. S3B), which inhibit pyruvate dehydrogenase and thereby limit conversion of pyruvate to acetyl-CoA for entry into the TCA cycle. Consistent with this restriction, pyruvate abundance accumulated in cortical neurons at 24 hpi (Fig. S3F). We and others have previously shown that in ZIKV-infected neurons, IRG1-derived itaconate inhibits succinate dehydrogenase (14), which drives accumulation of succinate and upstream α- ketoglutarate together with depletion of downstream fumarate and malate (Fig. S3G-J). Thus, our data suggest that ZIKV infection elicits distinct metabolic states in the two cell types: astrocytes adopted a more oxidative profile despite their typically glycolytic baseline, whereas neurons showed impaired mitochondrial carbon entry and accumulation of glycolytic and TCA intermediates. Given the increase in SIRT1 activity observed in infected astrocytes, these findings raise the possibility that SIRT1 drives this oxidative metabolic state and thereby supports conditions favorable for viral replication.

### SIRT1 activity sustains orthoflavivirus replication in astrocytes and exacerbates pathogenesis in vivo

We and others previously showed that suppressing oxidative phosphorylation while enhancing glycolysis establishes an antiviral metabolic state in neurons that limits flavivirus replication (14). Because SIRT1 activation in astrocytes was associated with a more oxidative metabolic profile, we next questioned whether SIRT1 activity influences viral replication in these cells. Primary human astrocytes were thus infected with ZIKV MR766 (MOI 0.01) in the presence of the SIRT1 inhibitor selisistat (10 μM), the SIRT1 activator SRT2104 (10 μM), or vehicle control, and infectious virus released into the culture supernatant was measured over 72 h (Fig. 4A-B). Selisistat reduced ZIKV titers relative to vehicle (Fig. 4A). Conversely, SRT2104 increased viral titers at 48 and 72 hpi relative to vehicle-treated cultures (Fig. 4B). These findings suggest that SIRT1 activity promotes ZIKV replication in primary human astrocytes.

**Figure 4.**
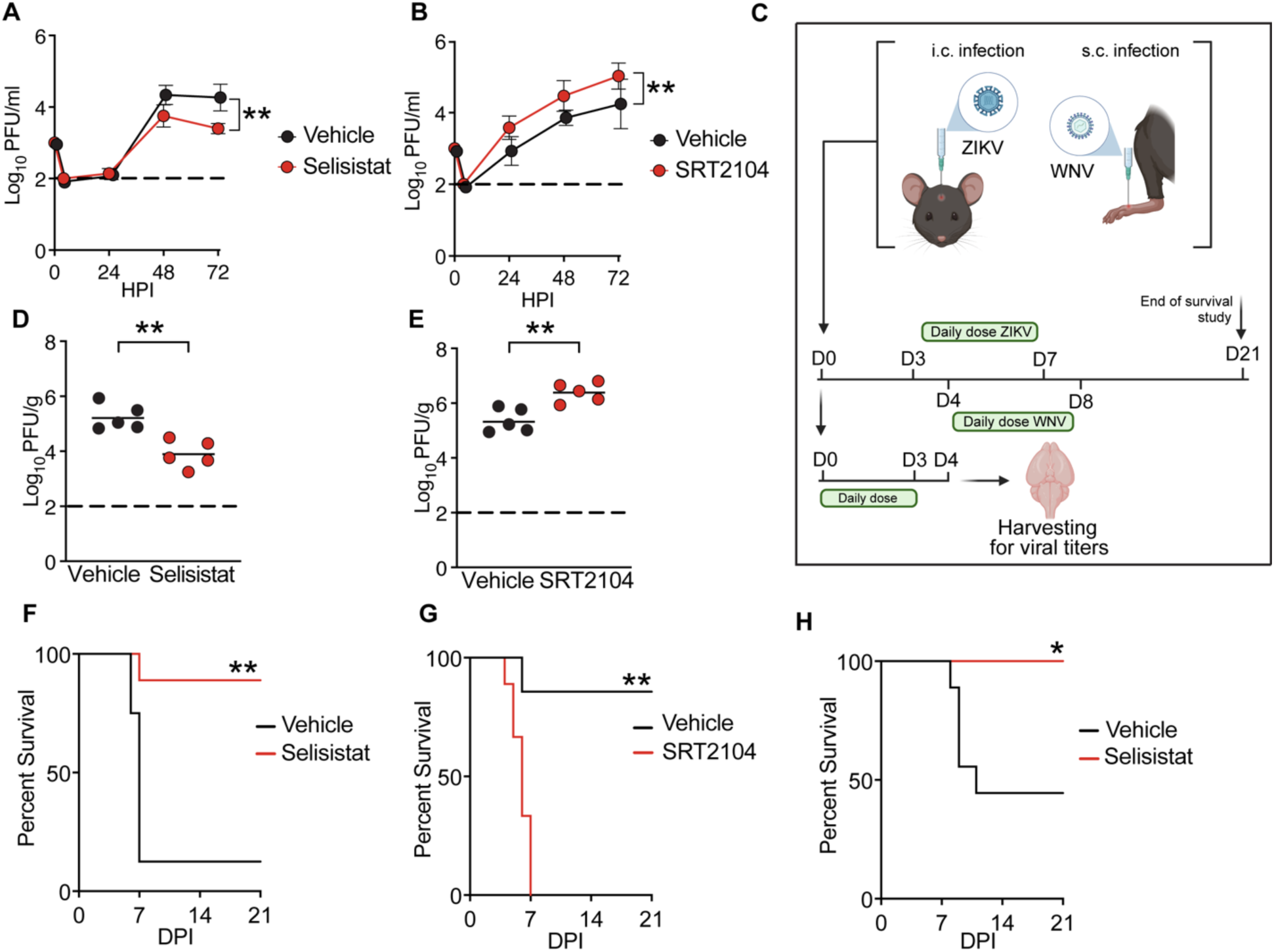
SIRT1 activity sustains orthoflavivirus replication in astrocytes and exacerbates pathogenesis in vivo. (A-B) Infectious ZIKV released into the culture supernatant of ZIKV-infected primary mouse astrocytes treated with selisistat (A), SRT2104 (B), or vehicle. n = 3 independent cultures /condition. Comparisons via two-way ANOVA. (C) Schematic of the in vivo treatment regimens used in this study. C57BL/6J mice were infected as indicated and treated intraperitoneally with daily injections of vehicle, selisistat (10 mg/kg), or SRT2104 (25 mg/kg). This panel was created with BioRender.com. (D-E) Brain viral burden in ZIKV-infected mice at 4 dpi following indicated treatments, quantified by plaque assay on whole-brain homogenates. n = 6 mice/group. Bars indicate the median. Comparisons via Mann-Whitney test. (F-H) Survival analysis of ZIKV-infected (F-G) or WNV-infected (H) mice receiving indicated treatments. n = 8 mice/group. Comparisons via log-rank test. Data in (A-B, D-E) represent mean values ± SEM. Horizontal lines in (D-E) indicate group medians. \**p* < 0.05, \*\**p* < 0.01.

We next asked whether pharmacologic modulation of SIRT1 similarly affected ZIKV replication in vivo. C57BL/6J mice were inoculated intracranially with ZIKV-MR766 and treated intraperitoneally with daily injections of vehicle, the SIRT1 inhibitor selisistat (10 mg/kg), or the SIRT1 activator SRT2104 (25 mg/kg) (Fig. 4C). We note that both of these compounds have been shown to cross the blood-brain barrier and impact SIRT1 signaling in the CNS following peripheral administration (44–47). Whole-brain homogenates were collected at 4 dpi, and infectious viral burden was quantified by plaque assay. Because we anticipated that SIRT1 inhibition and activation might exert opposing effects on disease severity, the selisistat experiment used a lethal inoculum of 10^4^ PFU, whereas the SRT2104 experiment used a sublethal inoculum of 10^1^ PFU. Selisistat treatment significantly reduced brain viral burden (Fig. 4D), while SRT2104 treatment increased brain viral burden (Fig. 4E). Thus, pharmacological inhibition and activation of SIRT1 produced reciprocal effects on ZIKV burden in the brain, consistent with the effects observed in primary human astrocytes and supporting a role for SIRT1 activity in promoting viral replication in vivo.

We next asked whether the effects of SIRT1 modulation on brain viral burden were accompanied by corresponding changes in disease outcome. C57BL/6J mice were infected intracranially with ZIKV-MR766 using either a lethal inoculum of 10^4^ PFU (for experiments using selisistat) or a sublethal inoculum of 10^1^ PFU (for experiments using SRT2104), as above. Drug treatment was initiated at 3 dpi and administered intraperitoneally once daily for 5 days using vehicle, selisistat (10 mg/kg), or SRT2104 (25 mg/kg), and survival was monitored through 21 dpi. Selisistat markedly improved survival following lethal ZIKV challenge, as vehicle-treated mice largely succumbed to infection by 7 dpi, whereas most selisistat-treated mice survived through the end of the observation period (Fig. 4F). Conversely, SRT2104 greatly exacerbated an otherwise largely sublethal challenge, with the majority of treated mice succumbing to infection by 6 dpi while most vehicle-treated animals survived through 21 dpi (Fig. 4G). We also examined whether SIRT1 inhibition altered disease caused by WNV. C57BL/6J mice were infected by footpad with 10^3^ PFU of WNV-Bird114 and treated intraperitoneally once daily for 5 days with vehicle or selisistat (10 mg/kg), beginning at 4 dpi. Selisistat treatment significantly reduced mortality in this infection model relative to vehicle treatment (Fig. 4H). Together, these experiments show that pharmacologic modulation of SIRT1 alters both viral replication and disease outcome following orthoflavivirus infection. Selisistat reduced ZIKV replication in primary human astrocytes and in the infected brain and improved survival following lethal ZIKV and WNV challenge. In contrast, SRT2104 increased ZIKV replication and worsened disease following an otherwise sublethal challenge. The reciprocal effects of SIRT1 inhibition and activation are consistent with a model in which SIRT1 enzymatic activity promotes efficient viral replication and contributes to CNS pathogenesis.

### The NAD+ salvage pathway sustains SIRT1-driven orthoflavivirus replication and in vivo pathogenesis

SIRT1 is an NAD+-dependent deacetylase, and the NAD+ salvage pathway helps maintain its cosubstrate availability through NAMPT-mediated conversion of NAM to NMN, followed by regeneration of NAD+ (Fig. 2A). We therefore asked whether pharmacologic manipulation of this pathway alters flavivirus replication. We targeted three distinct points in the NAD+ pathway: NAM, which at high concentrations can inhibit SIRT1 through product feedback; NMN, an NAD+ precursor that enters the salvage pathway downstream of NAMPT; and FK866, a small-molecule NAMPT inhibitor that reduces NAD+ synthesis. Primary human astrocytes were infected with ZIKV MR766 (MOI 0.01) in the presence of vehicle, NAM (10 mM), NMN (250 μM), or FK866 (100 nM), and infectious virus released into the culture supernatant was measured by plaque assay at 24, 48, and 72 hpi. NAM treatment reduced ZIKV replication across the time course relative to vehicle treatment (Fig. 5A), whereas NMN treatment increased ZIKV replication (Fig. 5B). FK866 treatment also reduced viral replication relative to vehicle treatment (Fig. 5C). Thus, interventions expected to limit NAD+-dependent SIRT1 activity reduced viral replication, whereas supplementation with an NAD+ precursor enhanced it. These effects parallel those observed with direct pharmacological modulation of SIRT1 and support a model in which NAD+ availability contributes to efficient ZIKV replication in astrocytes.

**Figure 5.**
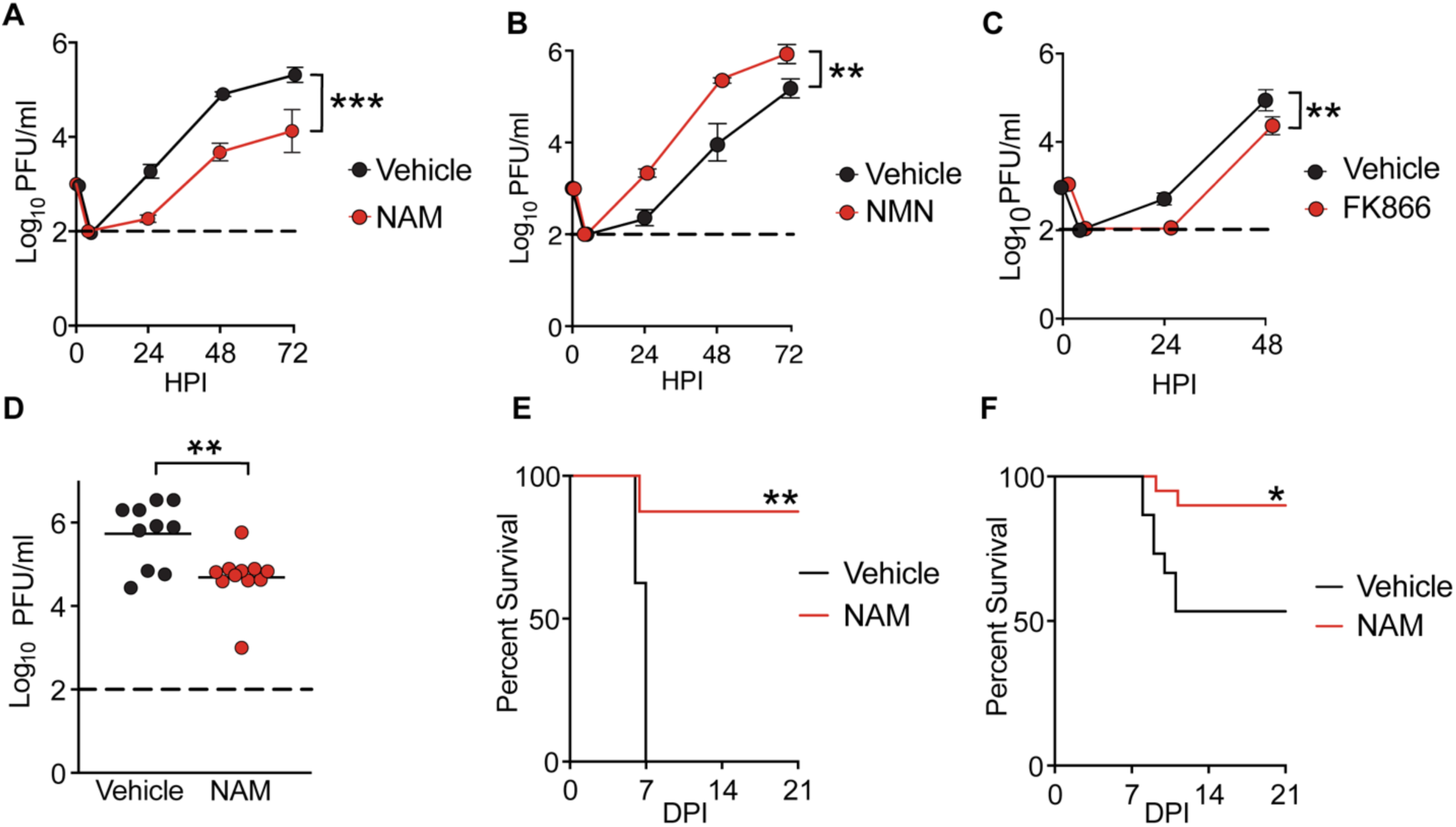
The NAD+ salvage pathway sustains SIRT1-driven orthoflavivirus replication and in vivo pathogenesis. (A–C) Infectious ZIKV released into the culture supernatant of ZIKV-infected primary mouse astrocytes treated with NAM (10 mM), NMN (250 μM), FK866 (100 nM) or vehicle. n = 4 independent cultures/condition. Comparisons via two-way ANOVA. (D) Brain viral burden in ZIKV-infected mice at 4 dpi following daily intraperitoneal treatment with vehicle or NAM (500 mg/kg), quantified by plaque assay on whole-brain homogenates. n = 10 mice/group. Comparisons via Mann-Whitney test. (E-F) Survival analysis of ZIKV-infected (E) or WNV-infected (F) mice treated with vehicle or NAM. n = 8 mice per group. Comparisons via log-rank test. Data in (A-C) represent mean values ± SEM. Horizontal lines in (D) indicate group medians. \**p* < 0.05, \*\**p* < 0.01, \*\*\**p* < 0.001.

We next asked whether NAM treatment would reproduce the effects of SIRT1 inhibition in vivo. C57BL/6J mice were infected either intracranially with ZIKV MR766 (10^4^ PFU) or by footpad with WNV-Bird114 (10^3^ PFU) and treated intraperitoneally with vehicle or NAM (500 mg/kg). Because NAM crosses the blood-brain barrier and enters the CNS following systemic delivery (48), we reasoned that peripheral administration would raise NAM availability in the infected brain. Daily NAM administration beginning on the day of infection significantly reduced brain ZIKV burden at 4 dpi (Fig. 5D). To assess disease outcome, separate cohorts of mice received five daily NAM injections from 3 through 7 dpi. NAM treatment improved survival following both lethal intracranial ZIKV challenge (Fig. 5E) and peripheral WNV infection (Fig. 5F). Across these measures, NAM produced effects similar to those observed with selisistat, including reduced brain viral burden and improved survival during ZIKV and WNV infection. These findings indicate that pharmacological manipulation of the NAD+ axis can influence orthoflavivirus replication and disease in vivo, consistent with a model in which NAD+ availability contributes to SIRT1- dependent pathogenesis.

## Discussion

SIRT1 is increasingly recognized as a central metabolic sensor that links cellular energetic state to transcriptional regulation through its dependence on NAD+ (49, 50). By coupling deacetylase activity to NAD+ availability, SIRT1 integrates metabolic information with programs controlling mitochondrial function, oxidative metabolism, stress resistance, and inflammatory signaling. Alterations in SIRT1 activity have been described in numerous viral infections, although whether these changes ultimately benefit the host or the pathogen appears highly context dependent and varies substantially across viruses and cell types (51–56). In the present study, we identify astrocytes as a previously unrecognized CNS cell population in which orthoflavivirus infection engages SIRT1 activity. This response was accompanied by transcriptional and metabolic changes consistent with enhanced NAD+ salvage and oxidative metabolism, and pharmacologic modulation of either SIRT1 or the NAD+ salvage pathway produced reciprocal effects on viral replication and disease severity. Together, these findings establish SIRT1 as an important component of astrocytic responses to orthoflavivirus infection and demonstrate that its activity influences the outcome of infection, although the upstream signals responsible for SIRT1 activation in astrocytes and the specific downstream mechanisms by which it affects viral replication require further investigation.

The established metabolic functions of SIRT1 provide some plausible hypotheses. During nutrient limitation and other forms of energetic stress, SIRT1 deacetylates PGC-1α to promote transcriptional programs supporting mitochondrial fatty acid oxidation and oxidative metabolism (57), while its regulation of FOXO proteins can promote resistance to oxidative damage (58). These activities help preserve mitochondrial function and metabolic flexibility when cellular demands exceed nutrient availability. Such adaptation could indirectly favor viral replication by sustaining ATP production and redox homeostasis while maintaining the mitochondrial carbon flux needed to generate amino acids, nucleotides, lipids, and other biosynthetic intermediates (59, 60). Preservation of host-cell metabolic capacity may be especially consequential in astrocytes, which can tolerate infection and support viral production over a relatively prolonged period (61). Thus, increased SIRT1 activity during infection may represent an active viral strategy to manipulate host metabolism in a manner favorable to viral replication. Alternatively, it may instead represent a host response to metabolic stress that incidentally drives the same result. Our findings are consistent with either model, although which SIRT1-dependent metabolic outputs may be required to promote viral replication remains to be determined.

A shift away from glycolytic metabolism may also impose important costs for astrocyte function during CNS infection. Astrocytes are intrinsically glycolytic cells, and glucose utilization is closely coupled to several of their homeostatic functions (62). Astrocytic glutamate uptake is an energy-intensive process that depends on glycolysis-derived ATP and is essential for limiting extracellular glutamate accumulation and excitotoxic neuronal injury (63, 64). Astrocytic glucose metabolism can also generate lactate that supports neighboring neurons under conditions of increased energetic demand, although the quantitative importance and directionality of lactate transfer remain debated (65). In addition, diversion of glucose through the pentose phosphate pathway supplies NADPH required for glutathione recycling and antioxidant defense (66), functions that may become particularly important during the oxidative stress associated with viral encephalitis. Suppression of glycolytic output in infected astrocytes could therefore impair neurotransmitter and ion homeostasis, reduce metabolic support for neurons, or weaken antioxidant capacity, thereby contributing to neuronal dysfunction independently of effects on viral replication (67). Our data do not establish that these functions are disrupted during infection, but they raise the possibility that the oxidative metabolic phenotype observed in astrocytes may simultaneously support viral production and compromise the protective functions these cells normally provide within the infected CNS.

The clinical effects of SIRT1 and NAD⁺ salvage modulation in our study suggest that this axis may be amenable to host-directed therapy during orthoflavivirus encephalitis (Fig. 6). Both approaches have precedent in other disease settings. Selisistat has been widely used in animal models of neurologic disease, as well as an exploratory clinical trial in patients with Huntington’s disease (68, 69). NAMPT inhibitors such as FK866/APO866 have likewise been developed as anticancer agents to exploit tumor dependence on NAD+ salvage; however, a phase 2 trial in cutaneous T-cell lymphoma showed limited activity and produced severe lymphocytopenia and thrombocytopenia, underscoring the potential toxicity of broadly disrupting NAD+ metabolism (70). Exogenous NAM administration may represent a more accessible alternative and has been used safely in human studies, including a phase 3 trial in which oral treatment reduced the incidence of new nonmelanoma skin cancers among high-risk adults (71). Notably, the acute nature of viral encephalitis may permit a comparatively brief period of metabolic modulation, potentially minimizing risks associated with long term treatments in more chronic disease states.

**Figure 6.**
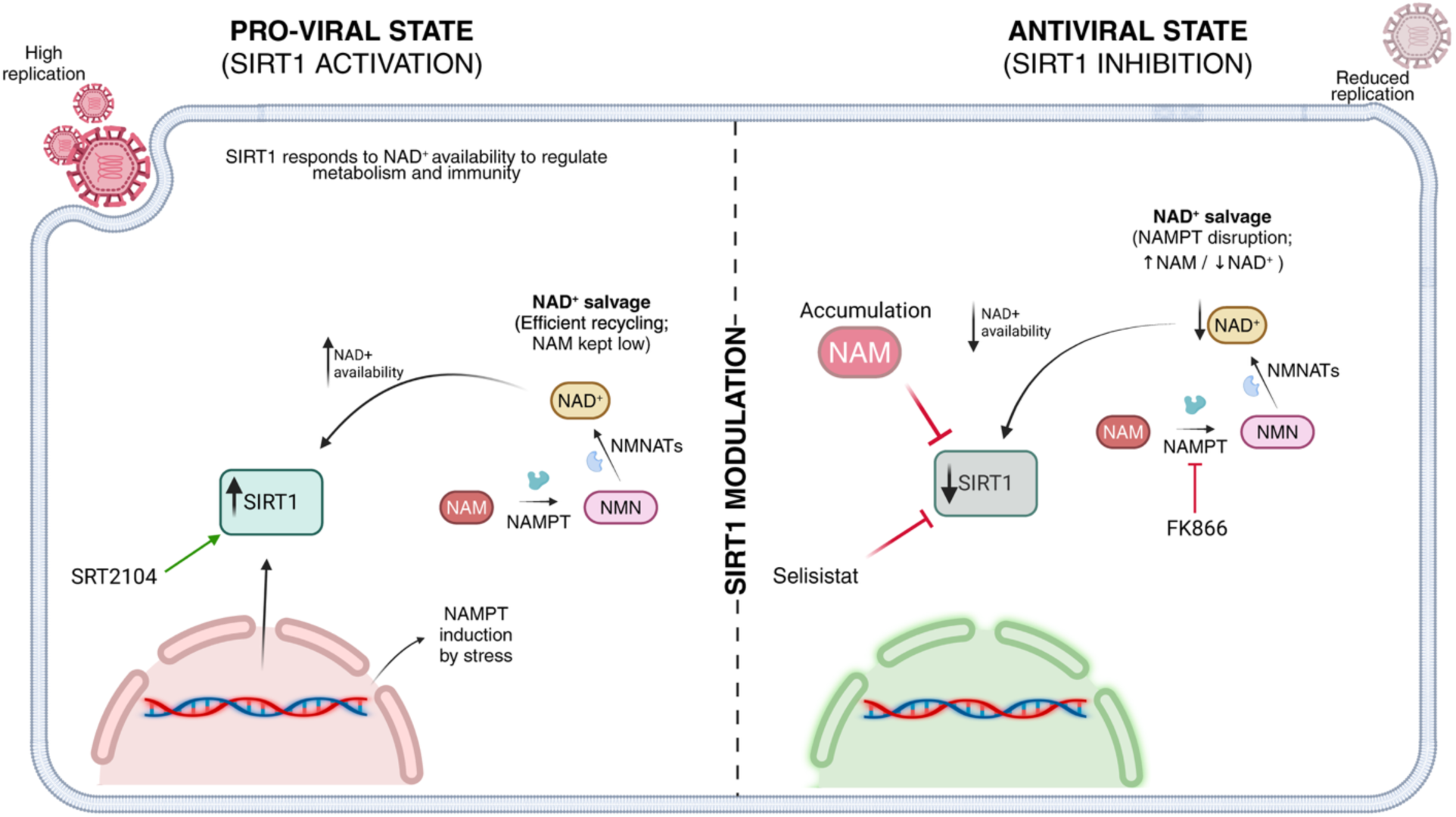
Proposed model of NAD⁺ salvage and SIRT1 modulation during orthoflavivirus infection. In infected astrocytes, induction of NAMPT and efficient recycling of nicotinamide (NAM) through nicotinamide mononucleotide (NMN) may preserve NAD⁺ availability, limit accumulation of the SIRT1-inhibitory product NAM, and sustain SIRT1 activity. This state, which is enhanced pharmacologically by the SIRT1 activator SRT2104, is associated with increased viral replication. Conversely, direct SIRT1 inhibition with selisistat, NAM accumulation, or disruption of NAD⁺ salvage through NAMPT inhibition with FK866 reduces SIRT1 activity and is associated with restricted viral replication. The schematic summarizes the pharmacologic relationships identified in this study and presents a proposed framework linking astrocyte NAD⁺ metabolism, SIRT1 activity, and orthoflavivirus replication. Figure created with BioRender.com.

Nevertheless, the central roles of SIRT1 and NAD+ in cellular homeostasis suggest that therapeutic approaches targeting these pathways specifically within astrocytes may achieve greater efficacy while minimizing the off-target effects and toxicity associated with systemic inhibition. Thus, future studies will need to carefully define the temporal and cell-specific contributions of the NAD+/SIRT1 axis during infection, identify the downstream metabolic functions most relevant to viral replication and tissue injury, and develop astrocyte-directed strategies that can modulate this pathway within a defined therapeutic window while preserving its essential functions in other CNS and peripheral cell populations.

## Methods

### Mice

C57BL/6J mice were bred and housed under specific pathogen-free conditions in Nelson Biological Laboratories at Rutgers University. All animals were congenic to the C57BL/6J background (Jackson Laboratories, #000664). Adult (>8-week-old) animals of both sexes were used in all studies, following protocols approved by the Rutgers University Institutional Animal Care and Use Committee (Protocol 201900016).

### Viruses and virologic assays

ZIKV-MR766 was provided by the World Reference Center for Emerging Viruses and Arboviruses (WRCEVA). WNV strain WN02-Bird 114 was generously provided by Dr. Bobby Brooke Herrera (Rutgers Robert Wood Johnson Medical School, New Brunswick, NJ, USA). Viral stocks were generated by infecting Vero cells (ATCC, #CCL-81) at a multiplicity of infection (MOI) of 0.01 and harvesting supernatants at 72 hours after infection. Viral titers of stocks were determined by plaque assay on Vero cells. Cells were maintained in DMEM (Corning #10-013-CV) supplemented with 10% heat-inactivated fetal bovine serum (FBS, Gemini Biosciences #100- 106), 1% Penicillin-Streptomycin-Glutamine (Gemini Biosciences #400-110), 1% Amphotericin B (Gemini Biosciences #400-104), 1% Non-Essential Amino Acids (Cytiva, #SH30238.01), and 1% HEPES (Cytiva, #SH30237.01). Plaque assay medium was composed of 1X EMEM (Lonza #12- 684F) supplemented with 2% heat-inactivated FBS (Gemini Biosciences #100-106), 1% Penicillin-Streptomycin-Glutamine (Gemini Biosciences, #400-110), 1% Amphotericin B (Gemini Biosciences #400-104), 1% Non-Essential Amino Acids (Cytiva, #SH30238.01), 1% HEPES (Cytiva, #SH30237.01), 0.75% Sodium Bicarbonate (VWR, #BDH9280), and 0.5% Methyl Cellulose (VWR, #K390). Plaque assays were performed by adding 100 μl of serially diluted sample for 1 hour at 37°C to 12-well plates containing 200,000 Vero cells per well. Plates were incubated with plaque assay medium at 37°C and 5% CO2 for 4 days. Plaque assays were developed by removal of overlay medium and staining/fixation with 10% neutral buffered formalin (VWR, #89370) and 0.25% crystal violet (VWR, #0528) for 20 to 30 min. Plates were washed repeatedly in H2O and allowed to dry before visible plaques were counted.

### Mouse infections and tissue harvesting

Isoflurane anesthesia was used for all procedures. Mice were inoculated intracranially (10 μl) with ZIKV-MR766 at 10^4^ PFU (lethal challenge) or 10^1^ PFU (sublethal challenge), or subcutaneously in a rear footpad (50 μl) with 10^3^ PFU of WNV-WN02-Bird114. Infected mice were monitored by a blinded operator daily for weight loss and presentation of clinical signs of disease, including hunched posture, ruffled fur, hindlimb weakness, and paresis. Mice reaching a moribund state or losing more than 20% of their initial body weight were euthanized, and the date of euthanasia was recorded for the purpose of survival studies. For brain harvest experiments, mice underwent cardiac perfusion with 30 ml of cold sterile 1X phosphate-buffered saline (PBS) at 4 days post-infection for ZIKV-infected mice or 9 days post-infection for WNV-infected mice. Extracted brains were weighed and homogenized with 1.0-mm diameter zirconia/silica beads (Biospec Products, #11079110z) in sterile PBS for plaque assays. Homogenization was performed in an Omni Beadrupter Elite for two sequential cycles of 20 s at a speed of 4 m/s.

### In vivo drug treatments

Selisistat (EX-527, MedChemExpress, #HY-15452) and SRT2104 (Selleckchem, #S7792) were reconstituted in 10% DMSO and 90% polyethylene glycol 400 (PEG 400). Nicotinamide (NAM, Sigma-Aldrich, #N0636) was dissolved in sterile PBS. All drugs were administered by intraperitoneal injection in a 200 μl volume. For ZIKV brain viral burden experiments, mice received daily injections of selisistat (10 mg/kg), SRT2104 (25 mg/kg), NAM (500 mg/kg), or vehicle from day 0 through day 3 post-infection, and brains were harvested at 4 days post-infection. For ZIKV survival experiments, treatment with selisistat (10 mg/kg), SRT2104 (25 mg/kg), NAM (500 mg/kg), or vehicle was initiated at 3 days post-infection and continued daily for 5 consecutive days. For WNV survival experiments, treatment with selisistat (10 mg/kg), SRT2104 (25 mg/kg), NAM (500 mg/kg), or vehicle was initiated at 4 days post-infection and continued daily for 5 consecutive days.

### Primary cell culture

Primary mouse cortical astrocytes were generated from P1 to P3 C57BL/6J mouse pups as previously described (72). Tissues were dissociated with the Neural Dissociation Kit (T) according to the manufacturer’s instructions (Miltenyi, #130-093-231). Astrocytes were expanded in AM-a medium (ScienCell, #1831) supplemented with 10% FBS (Gemini Biosciences, #100-106) in fibronectin-coated cell culture flasks and seeded into plates coated with Poly-L-Lysine (20 μg/ml, Sigma-Aldrich, #9155) before experiments. Pooled tissues from both male and female pups were used for all primary mouse astrocyte cultures. Primary human cortical astrocytes (ScienCell, #1800) were cultured in astrocyte medium (ScienCell, #1801) supplemented with 2% heat-inactivated FBS (ScienCell, #0010), astrocyte growth supplement (ScienCell, #1852), and penicillin/streptomycin cocktail (ScienCell, #0503). Human astrocytes were maintained in poly-L-lysine coated flasks and used within three passages.

### In vitro infections and drug treatments

Primary mouse cortical astrocytes and primary human cortical astrocytes were infected with ZIKV- MR766 at a multiplicity of infection (MOI) of 0.01. For viral growth curve experiments, supernatants from infected primary human astrocytes were collected at 24, 48, and 72 hours post-infection for plaque assay. For pharmacological treatments, the following compounds were added at the time of infection at the indicated final concentrations: selisistat (EX-527, 10 μM, MedChemExpress, #HY-15452), SRT2104 (10 μM, Selleckchem, #S7792), nicotinamide (NAM, 10 mM, Sigma-Aldrich, #N0636), β-nicotinamide mononucleotide (NMN, 250 μM, MedChemExpress, #HY-F0004), or FK866 (100 nM, MedChemExpress, #HY-50876). DMSO was used as vehicle control at 0.1% final concentration.

### Magnetic activated cell sorting (MACS)

MACS was performed with commercial kits according to manufacturer’s instructions. Mouse brains were dissociated with the Adult Brain Dissociation Kit (Miltenyi, #130-107-677), and isolations were performed with microbead kits designed to isolate astrocytes (Anti-ACSA-2 MicroBead Kit, Miltenyi, #130-097-678), neurons (Neuron Isolation Kit, mouse, Miltenyi, #130- 115-389), and myeloid cells (CD11b MicroBeads, Miltenyi, #130-126-725). Brains were harvested at 4 days post-infection for ZIKV-infected mice and at 9 days post-infection for WNV-infected mice.

### SIRT1 deacetylase activity assay

SIRT1 deacetylase activity was measured using the SensoLyte Green SIRT1 Assay Kit (AnaSpec, #AS-72156) according to the manufacturer’s instructions. For in vivo experiments, astrocytes, neurons, and myeloid cells isolated by MACS from mock-infected, ZIKV-infected (4 dpi), or WNV-infected (9 dpi) C57BL/6J mouse brains were used. For in vitro experiments, primary human cortical astrocytes were collected at 72 hours after mock infection or ZIKV-MR766 infection (MOI 0.01). Cells were lysed in the kit-provided assay buffer, and protein concentration was determined by BCA assay (ThermoFisher Scientific, #23227). Equal amounts of protein were incubated with the fluorogenic SIRT1 substrate in a black-walled 96-well plate at 37°C, and fluorescence was measured on a SpectraMax iD3 plate reader (Molecular Devices, San Jose, CA) at excitation 490 nm and emission 520 nm.

### Metabolite quantification by liquid chromatography-mass spectrometry

Whole brain tissue from ZIKV-infected and mock-infected mice at 4 days post-infection, and primary human cortical astrocytes at 48 hours after mock or ZIKV-MR766 infection (MOI 0.01), were flash-frozen in liquid nitrogen at the time of harvest and submitted to the Metabolomics Shared Resource Facility at Rutgers Robert Wood Johnson Medical School (New Brunswick, NJ). Metabolite extraction, hydrophilic interaction liquid chromatography (HILIC), and high-resolution mass spectrometry analysis were performed by the core. Peak areas for metabolites of interest, including NAD+, NADH, NAM, glucose, pyruvate, lactate, acetyl-CoA, aspartate, and glutamate, were extracted from the untargeted dataset. Metabolite levels were normalized to tissue weight for whole brain samples and to cell number for cultured astrocyte samples. Metabolomic analysis in neurons was performed using a previously published dataset which was generated as described (14).

### Nicotinamide ELISA

NAM concentrations in some samples were measured using the Nicotinamide (NAM) ELISA Kit (MyBioSource, #MBS7269283) according to the manufacturer’s instructions. Samples included MACS-sorted astrocytes from ZIKV-infected mouse brains at 4 days post-infection, WNV-infected whole brain homogenates at 9 days post-infection, primary human cortical astrocytes at 72 hours after mock or ZIKV-MR766 infection (MOI 0.01), and primary mouse cortical astrocytes at 72 hours after mock, ZIKV-MR766, or WNV-WN02-Bird114 infection (MOI 0.01). Cells and brain tissue were lysed in ice-cold PBS by sonication, centrifuged at 14,000 × g for 10 minutes at 4°C, and clarified supernatants were used for the assay. Absorbance was read at 450 nm on a SpectraMax iD3 plate reader (Molecular Devices, San Jose, CA). NAM concentrations were normalized to total protein, quantified in parallel by BCA assay (ThermoFisher Scientific, #23227).

### Mitochondrial respiration and glycolysis assays

Oxygen consumption rate (OCR) and extracellular acidification rate (ECAR) were measured in primary mouse cortical astrocytes using the Extracellular Oxygen Consumption Assay Kit (Abcam, #ab197243) and the Glycolysis Assay (Extracellular Acidification) Kit (Abcam, #ab197244), respectively, according to the manufacturer’s instructions. Primary mouse astrocytes were seeded at equal density in black-walled 96-well plates and infected with ZIKV-MR766 (MOI 0.01) or mock-infected. At 72 hours post-infection, cells were equilibrated with assay medium containing the oxygen-sensing or pH-sensing fluorescent probe and overlaid with mineral oil.

Kinetic fluorescence was recorded at 37°C on a SpectraMax iD3 plate reader (Molecular Devices, San Jose, CA) by time-resolved fluorescence (OCR at excitation 380 nm and emission 650 nm, ECAR at excitation 380 nm and emission 615 nm). OCR and ECAR rates were calculated from the linear slope of the fluorescence-time curve.

### RNA sequencing analysis

Primary mouse cortical astrocytes were infected with ZIKV-MR766 (MOI 0.01) or mock-infected and collected at 48 and 72 hours post-infection, with three biological replicates per condition. Total RNA was extracted and submitted to Azenta Life Sciences for library preparation, next-generation sequencing, and initial processing as previously described (73, 74). Sequence reads were quality-checked, adapter-trimmed, aligned to the mouse reference genome, and quantified to generate a gene-level count matrix. The gene-level count matrix delivered by Azenta was imported into R (v4.5.1) for downstream analysis. Counts were filtered to retain genes with a row sum of at least 10 reads across all samples, duplicate gene symbols were collapsed by retaining the highest-expressed row, and Ensembl IDs were mapped to gene symbols using org.Mm.eg.db. Differential expression was tested with DESeq2 (v1.48.2) using a design of ∼ condition_timepoint with three levels (Mock 48 hpi, ZIKV 48 hpi, ZIKV 72 hpi) and Mock 48 hpi set as the reference. Contrasts were extracted for ZIKV 48 hpi vs Mock 48 hpi and ZIKV 72 hpi vs Mock 48 hpi, with significance assessed at alpha = 0.05 (Benjamini-Hochberg adjusted).

For comparison with cortical neuron transcriptional responses, microarray data from primary mouse cortical neurons infected with ZIKV-MR766 (MOI 0.1) or WNV-TX 2002-HC (MOI 0.001) at 24 hpi (NCBI Gene Expression Omnibus accession GSE122121) were downloaded from the NCBI Gene Expression Omnibus (14). Values associated with wild-type samples were extracted, and differential expression was assessed using limma with lmFit followed by empirical Bayes moderation (eBayes) (75). The moderated t-statistic served as the input metric for downstream transcription factor activity inference.

Transcription factor activity was inferred using the univariate linear model method (run_ulm) in decoupleR with the DoRothEA mouse regulon collection at confidence levels A through C (273 transcription factors) restricted to regulons with at least 15 target genes (76–78). The DESeq2 Wald statistic (log2FoldChange / lfcSE) served as the ranking metric for the astrocyte data, and the moderated t-statistic served as the ranking metric for the neuron data. SIRT1 substrates analyzed comprised the activated arm (Foxo1, Foxo3, Foxo4, Arntl, Nfe2l2, Nr1h3, Epas1, Hsf1) and the repressed arm (E2f1, Myc, Srebf1). FDR was thresholded at 0.05. Per-gene log2 fold changes within each SIRT1 substrate regulon were compared between 48 and 72 hpi by paired Wilcoxon signed-rank test.

Pre-ranked gene set enrichment analysis was performed using fgsea (v1.34.20) (79) using the 72 hpi Wald statistic as the ranking metric (40). Gene sets corresponded to the DoRothEA mouse regulon target gene lists for Foxo1 (43 targets), Foxo3 (39 targets), E2f1 (307 targets), and Myc (363 targets). Analysis parameters were eps = 0.0, minSize = 5, and set.seed = 42. Mode of regulation weighting was not applied.

Canonical pathway and Upstream Regulator analyses were performed in QIAGEN Ingenuity Pathway Analysis (IPA, Application Build 9.0, Content version 159584291). The DESeq2 output (genes with Benjamini-Hochberg adjusted p-value less than 0.05) was uploaded to IPA, and pathway and regulator activation states were inferred from the IPA-calculated z-scores.

## Statistical analysis

Statistical analyses were performed using GraphPad Prism v10 (GraphPad Software, San Diego, CA) and R (v4.5.1). Welch’s two-tailed t-test was used for pairwise comparisons of metabolite levels, enzymatic activities, and DNA-binding activities between mock-infected and infected groups. Mann-Whitney tests were used for comparisons of brain viral titers across treatment groups. Two-way analysis of variance (ANOVA) was used for time-course plaque assay comparisons across timepoints and treatments. Log-rank tests were used for survival curve comparisons. Paired Wilcoxon signed-rank tests were used for per-gene log2 fold change comparisons across timepoints within transcription factor regulons. False discovery rate was controlled by the Benjamini-Hochberg procedure for high-dimensional analyses. P < 0.05 was considered statistically significant unless otherwise specified. All data points represent biological replicates consisting of distinct mice or independent cultures derived from distinct mice unless otherwise noted. No data points were excluded from any analysis.

## Acknowledgements

The authors thank Dr. Xiaoyang Su and the Rutgers Cancer Institute Metabolomics Shared Resource, who are supported, in part, by NCI-CCSG P30CA072720-6852. This work was supported by R01 NS120895 (to B.P.D). J.P.A. was supported by a National Science Foundation Graduate Research Fellowship.

## Author Contributions

Conceptualization: JPA and BPD; Investigation: JPA, DA, ML, CA, and BPD; Analysis: JPA, DA, ML, and BPD; Writing – original draft: JPA and BPD; Writing – reviewing and editing: JPA and BPD; Supervision: CA and BPD; Funding Acquisition: BPD.

## Data Availability

Transcriptomic analysis in neurons was performed using a previously published dataset (GEO accession number: GSE122121). Transcriptomic analysis in astrocytes was performed using an unpublished dataset that will be made available upon publication.

## References

1. Fajardo A, Cristina J, Moreno P. Emergence and Spreading Potential of Zika Virus. Front Microbiol. 2016;7:1667.

2. Singh P, Khatib MN, Ballal S, Kaur M, Nathiya D, Sharma S, et al. West Nile Virus in a changing climate: epidemiology, pathology, advances in diagnosis and treatment, vaccine designing and control strategies, emerging public health challenges - a comprehensive review. Emerg Microbes Infect. 2025;14(1):2437244.

3. Faria NR, Quick J, Claro IM, Theze J, de Jesus JG, Giovanetti M, et al. Establishment and cryptic transmission of Zika virus in Brazil and the Americas. Nature. 2017;546(7658):406- 10.

4. Daniels BP, Jujjavarapu H, Durrant DM, Williams JL, Green RR, White JP, et al. Regional astrocyte IFN signaling restricts pathogenesis during neurotropic viral infection. J Clin Invest. 2017;127(3):843–56.

5. Telikani Z, Monson EA, Hofer MJ, Helbig KJ. Antiviral response within different cell types of the CNS. Front Immunol. 2022;13:1044721.

6. Lindman M, Angel JP, Estevez I, Chang NP, Chou TW, McCourt M, et al. RIPK3 promotes brain region-specific interferon signaling and restriction of tick-borne flavivirus infection. PLoS Pathog. 2023;19(11):e1011813.

7. Lang R, Li H, Luo X, Liu C, Zhang Y, Guo S, et al. Expression and mechanisms of interferon-stimulated genes in viral infection of the central nervous system (CNS) and neurological diseases. Front Immunol. 2022;13:1008072.

8. Jordan TX, Randall G. Flavivirus modulation of cellular metabolism. Curr Opin Virol. 2016;19:7–10.

9. Darweesh M, Mohammadi S, Rahmati M, Al-Hamadani M, Al-Harrasi A. Metabolic reprogramming in viral infections: the interplay of glucose metabolism and immune responses. Front Immunol. 2025;16:1578202.

10. Fontaine KA, Sanchez EL, Camarda R, Lagunoff M. Dengue virus induces and requires glycolysis for optimal replication. J Virol. 2015;89(4):2358–66.

11. Singh S, Singh PK, Suhail H, Arumugaswami V, Pellett PE, Giri S, et al. AMP-Activated Protein Kinase Restricts Zika Virus Replication in Endothelial Cells by Potentiating Innate Antiviral Responses and Inhibiting Glycolysis. J Immunol. 2020;204(7):1810–24.

12. Xu L, Li M, Zhang J, Li D, Tao J, Zhang F, et al. Metabolomic landscape of macrophage discloses an anabolic signature of dengue virus infection and antibody-dependent enhancement of viral infection. PLoS Negl Trop Dis. 2024;18(2):e0011923.

13. Tiwari SK, Dang J, Qin Y, Lichinchi G, Bansal V, Rana TM. Zika virus infection reprograms global transcription of host cells to allow sustained infection. Emerg Microbes Infect. 2017;6(4):e24.

14. Daniels BP, Kofman SB, Smith JR, Norris GT, Snyder AG, Kolb JP, et al. The Nucleotide Sensor ZBP1 and Kinase RIPK3 Induce the Enzyme IRG1 to Promote an Antiviral Metabolic State in Neurons. Immunity. 2019;50(1):64–76 e4.

15. Ledur PF, Karmirian K, Pedrosa C, Souza LRQ, Assis-de-Lemos G, Martins TM, et al. Zika virus infection leads to mitochondrial failure, oxidative stress and DNA damage in human iPSC-derived astrocytes. Sci Rep. 2020;10(1):1218.

16. Braz-de-Melo HA, Lago FG, Correa R, Santos IO, Bellozi PMQ, Almeida RDN, et al. Zika Virus Reprograms Microglial Mitochondrial Metabolism to Support Immune Activation and Viral Replication: Omega-3 DHA Counteracts Neuroinflammation and Viral Persistence. Mol Neurobiol. 2026;63(1):341.

17. de Farias IS, Ribeiro G, Noronha IH, Lucena VWL, Peron JPS, Moraes-Vieira PM, et al. Caspase-1/11 controls Zika virus replication in astrocytes by inhibiting glycolytic metabolism. FEBS J. 2025;292(12):3113–28.

18. Bonvento G, Bolanos JP. Astrocyte-neuron metabolic cooperation shapes brain activity. Cell Metab. 2021;33(8):1546–64.

19. Kabiraj P, Grund EM, Clarkson BDS, Johnson RK, LaFrance-Corey RG, Lucchinetti CF, et al. Teriflunomide shifts the astrocytic bioenergetic profile from oxidative metabolism to glycolysis and attenuates TNFalpha-induced inflammatory responses. Sci Rep. 2022;12(1):3049.

20. Lindqvist R, Mundt F, Gilthorpe JD, Wolfel S, Gekara NO, Kroger A, et al. Fast type I interferon response protects astrocytes from flavivirus infection and virus-induced cytopathic effects. J Neuroinflammation. 2016;13(1):277.

21. Stefanik M, Formanova P, Bily T, Vancova M, Eyer L, Palus M, et al. Characterisation of Zika virus infection in primary human astrocytes. BMC Neurosci. 2018;19(1):5.

22. Corcoran SE, O’Neill LA. HIF1alpha and metabolic reprogramming in inflammation. J Clin Invest. 2016;126(10):3699–707.

23. Brunet A, Sweeney LB, Sturgill JF, Chua KF, Greer PL, Lin Y, et al. Stress-dependent regulation of FOXO transcription factors by the SIRT1 deacetylase. Science. 2004;303(5666):2011–5.

24. Lim JH, Lee YM, Chun YS, Chen J, Kim JE, Park JW. Sirtuin 1 modulates cellular responses to hypoxia by deacetylating hypoxia-inducible factor 1alpha. Mol Cell. 2010;38(6):864–78.

25. Vaziri H, Dessain SK, Ng Eaton E, Imai SI, Frye RA, Pandita TK, et al. hSIR2(SIRT1) functions as an NAD-dependent p53 deacetylase. Cell. 2001;107(2):149–59.

26. Yeung F, Hoberg JE, Ramsey CS, Keller MD, Jones DR, Frye RA, et al. Modulation of NF-kappaB-dependent transcription and cell survival by the SIRT1 deacetylase. EMBO J. 2004;23(12):2369–80.

27. Qin Z, Fang X, Sun W, Ma Z, Dai T, Wang S, et al. Deactylation by SIRT1 enables liquid-liquid phase separation of IRF3/IRF7 in innate antiviral immunity. Nat Immunol. 2022;23(8):1193–207.

28. Yu SS, Tang RC, Zhang A, Geng S, Yu H, Zhang Y, et al. Deacetylase Sirtuin 1 mitigates type I IFN- and type II IFN-induced signaling and antiviral immunity. J Virol. 2024;98(3):e0008824.

29. Covarrubias AJ, Perrone R, Grozio A, Verdin E. NAD(+) metabolism and its roles in cellular processes during ageing. Nat Rev Mol Cell Biol. 2021;22(2):119–41.

30. Garten A, Petzold S, Korner A, Imai S, Kiess W. Nampt: linking NAD biology, metabolism and cancer. Trends Endocrinol Metab. 2009;20(3):130–8.

31. Avalos JL, Bever KM, Wolberger C. Mechanism of sirtuin inhibition by nicotinamide: altering the NAD(+) cosubstrate specificity of a Sir2 enzyme. Mol Cell. 2005;17(6):855–68.

32. Harlan BA, Pehar M, Sharma DR, Beeson G, Beeson CC, Vargas MR. Enhancing NAD+ Salvage Pathway Reverts the Toxicity of Primary Astrocytes Expressing Amyotrophic Lateral Sclerosis-linked Mutant Superoxide Dismutase 1 (SOD1). J Biol Chem. 2016;291(20):10836–46.

33. Pang H, Jiang Y, Li J, Wang Y, Nie M, Xiao N, et al. Aberrant NAD(+) metabolism underlies Zika virus-induced microcephaly. Nat Metab. 2021;3(8):1109–24.

34. Feige JN, Auwerx J. Transcriptional targets of sirtuins in the coordination of mammalian physiology. Curr Opin Cell Biol. 2008;20(3):303–9.

35. Jing H, Lin H. Sirtuins in epigenetic regulation. Chem Rev. 2015;115(6):2350–75.

36. Daitoku H, Hatta M, Matsuzaki H, Aratani S, Ohshima T, Miyagishi M, et al. Silent information regulator 2 potentiates Foxo1-mediated transcription through its deacetylase activity. Proc Natl Acad Sci U S A. 2004;101(27):10042–7.

37. van der Horst A, Tertoolen LG, de Vries-Smits LM, Frye RA, Medema RH, Burgering BM. FOXO4 is acetylated upon peroxide stress and deacetylated by the longevity protein hSir2(SIRT1). J Biol Chem. 2004;279(28):28873–9.

38. Xue F, Huang JW, Ding PY, Zang HG, Kou ZJ, Li T, et al. Nrf2/antioxidant defense pathway is involved in the neuroprotective effects of Sirt1 against focal cerebral ischemia in rats after hyperbaric oxygen preconditioning. Behav Brain Res. 2016;309:1–8.

39. Zhou Y, Peng L, Li Y, Zhao Y. Silent information regulator 1 ameliorates oxidative stress injury via PGC-1alpha/PPARgamma-Nrf2 pathway after ischemic stroke in rat. Brain Res Bull. 2022;178:37–48.

40. Hariharan N, Maejima Y, Nakae J, Paik J, Depinho RA, Sadoshima J. Deacetylation of FoxO by Sirt1 Plays an Essential Role in Mediating Starvation-Induced Autophagy in Cardiac Myocytes. Circ Res. 2010;107(12):1470–82.

41. Gerhart-Hines Z, Rodgers JT, Bare O, Lerin C, Kim SH, Mostoslavsky R, et al. Metabolic control of muscle mitochondrial function and fatty acid oxidation through SIRT1/PGC-1alpha. EMBO J. 2007;26(7):1913–23.

42. Estevez I, Buckley BD, Lindman M, Panzera N, Chou TW, McCourt M, et al. The kinase RIPK3 promotes neuronal survival by suppressing excitatory neurotransmission during central nervous system viral infection. Immunity. 2025;58(3):666–82 e6.

43. Angel JP, Daniels BP. Paradoxical roles for programmed cell death signaling during viral infection of the central nervous system. Curr Opin Neurobiol. 2022;77:102629.

44. Chang N, Li J, Lin S, Zhang J, Zeng W, Ma G, et al. Emerging roles of SIRT1 activator, SRT2104, in disease treatment. Sci Rep. 2024;14(1):5521.

45. Smith MR, Syed A, Lukacsovich T, Purcell J, Barbaro BA, Worthge SA, et al. A potent and selective Sirtuin 1 inhibitor alleviates pathology in multiple animal and cell models of Huntington’s disease. Hum Mol Genet. 2014;23(11):2995–3007.

46. Jiang M, Zheng J, Peng Q, Hou Z, Zhang J, Mori S, et al. Sirtuin 1 activator SRT2104 protects Huntington’s disease mice. Ann Clin Transl Neurol. 2014;1(12):1047–52.

47. Wei X, Chen S, Liu D, Li J, Deng Q, Wang Y, et al. The SIRT1 activator SRT2104 mitigates hypoxia-induced white matter injury in neonatal mice. Brain Res. 2025;1866:149917.

48. Fricker RA, Green EL, Jenkins SI, Griffin SM. The Influence of Nicotinamide on Health and Disease in the Central Nervous System. Int J Tryptophan Res. 2018;11:1178646918776658.

49. Thapa R, Moglad E, Afzal M, Gupta G, Bhat AA, Hassan Almalki W, et al. The role of sirtuin 1 in ageing and neurodegenerative disease: A molecular perspective. Ageing Res Rev. 2024;102:102545.

50. Wu QJ, Zhang TN, Chen HH, Yu XF, Lv JL, Liu YY, et al. The sirtuin family in health and disease. Signal Transduct Target Ther. 2022;7(1):402.

51. Wang Y, Li H, Huang X, Huang Y, Lv M, Tang H, et al. NAD+ Suppresses EV-D68 Infection by Enhancing Anti-Viral Effect of SIRT1. Viruses. 2025;17(2).

52. Wang J, Qin X, Huang Y, Zhang G, Liu Y, Cui Y, et al. Sirt1 Negatively Regulates Cellular Antiviral Responses by Preventing the Cytoplasmic Translocation of Interferon-Inducible Protein 16 in Human Cells. J Virol. 2023;97(2):e0197522.

53. Liu Q, He Q, Tao X, Yu P, Liu S, Xie Y, et al. Resveratrol inhibits rabies virus infection in N2a cells by activating the SIRT1/Nrf2/HO-1 pathway. Heliyon. 2024;10(17):e36494.

54. Chen D, Zhao G, Zhou J, Sun P, He S, Lv C, et al. Delactylation of viral proteins by SIRT1 suppresses influenza A virus replication. mBio. 2026;17(4):e0248925.

55. Yan Y, Ren X, Xiao Y, Li F, Guo J, Ji K, et al. The SIRT1-Mediated p53 Deacetylation Pathway Modulates Apoptosis and Promotes Viral Replication in MVC-Infected Cells. Pathogens. 2026;15(3).

56. Hajra D, Chakravortty D. Sirtuins as modulators of infection outcomes in the battle of host-pathogen dynamics. Phys Life Rev. 2025;53:225–35.

57. Chen J, Liu B, Yao X, Yang X, Sun J, Yi J, et al. AMPK/SIRT1/PGC-1alpha Signaling Pathway: Molecular Mechanisms and Targeted Strategies From Energy Homeostasis Regulation to Disease Therapy. CNS Neurosci Ther. 2025;31(11):e70657.

58. Guan G, Chen Y, Dong Y. Unraveling the AMPK-SIRT1-FOXO Pathway: The In-Depth Analysis and Breakthrough Prospects of Oxidative Stress-Induced Diseases. Antioxidants (Basel). 2025;14(1).

59. Kayesh MEH, Kohara M, Tsukiyama-Kohara K. Effects of oxidative stress on viral infections: an overview. Npj Viruses. 2025;3(1):27.

60. Palmer CS. Innate metabolic responses against viral infections. Nat Metab. 2022;4(10):1245–59.

61. Jorgacevski J, Potokar M. Immune Functions of Astrocytes in Viral Neuroinfections. Int J Mol Sci. 2023;24(4).

62. Beard E, Lengacher S, Dias S, Magistretti PJ, Finsterwald C. Astrocytes as Key Regulators of Brain Energy Metabolism: New Therapeutic Perspectives. Front Physiol. 2021;12:825816.

63. Silva-Hucha S, Hernandez RG, Baena-Lopez D, Fernandez de Sevilla ME, Paradas C, Morcuende S. Excitotoxicity in amyotrophic lateral sclerosis: a key pathogenic mechanism. Brain Commun. 2026;8(2):fcag098.

64. Zhang YM, Qi YB, Gao YN, Chen WG, Zhou T, Zang Y, et al. Astrocyte metabolism and signaling pathways in the CNS. Front Neurosci. 2023;17:1217451.

65. Kim Y, Dube SE, Park CB. Brain energy homeostasis: the evolution of the astrocyte-neuron lactate shuttle hypothesis. Korean J Physiol Pharmacol. 2025;29(1):1–8.

66. TeSlaa T, Ralser M, Fan J, Rabinowitz JD. The pentose phosphate pathway in health and disease. Nat Metab. 2023;5(8):1275–89.

67. Xiong J, Ge X, Pan D, Zhu Y, Zhou Y, Gao Y, et al. Metabolic reprogramming in astrocytes prevents neuronal death through a UCHL1/PFKFB3/H4K8la positive feedback loop. Cell Death Differ. 2025;32(7):1214–30.

68. Hong JY, Lin H. Sirtuin Modulators in Cellular and Animal Models of Human Diseases. Front Pharmacol. 2021;12:735044.

69. Sussmuth SD, Haider S, Landwehrmeyer GB, Farmer R, Frost C, Tripepi G, et al. An exploratory double-blind, randomized clinical trial with selisistat, a SirT1 inhibitor, in patients with Huntington’s disease. Br J Clin Pharmacol. 2015;79(3):465–76.

70. Goldinger SM, Gobbi Bischof S, Fink-Puches R, Klemke CD, Dreno B, Bagot M, et al. Efficacy and Safety of APO866 in Patients With Refractory or Relapsed Cutaneous T-Cell Lymphoma: A Phase 2 Clinical Trial. JAMA Dermatol. 2016;152(7):837–9.

71. Chen AC, Martin AJ, Choy B, Fernandez-Penas P, Dalziell RA, McKenzie CA, et al. A Phase 3 Randomized Trial of Nicotinamide for Skin-Cancer Chemoprevention. N Engl J Med. 2015;373(17):1618–26.

72. Lindman M, Estevez I, Marmut E, DaPrano EM, Chou TW, Newman K, et al. Astrocytic RIPK3 exerts protective anti-inflammatory activity in mice with viral encephalitis by transcriptional induction of serpins. Sci Signal. 2025;18(895):eadq6422.

73. Chou TW, McCourt M, Marmut E, Karuppusamy V, Lindman M, Estevez I, et al. Inapparent maternal ZIKV infection impacts fetal brain development and postnatal behavior. PLoS Pathog. 2026;22(1):e1013850.

74. Chang NP, DaPrano EM, Lindman M, Estevez I, Chou TW, Evans WR, et al. Neuronal DAMPs exacerbate neurodegeneration via astrocytic RIPK3 signaling. JCI Insight. 2024;9(11).

75. Smyth GK. Linear models and empirical bayes methods for assessing differential expression in microarray experiments. Stat Appl Genet Mol Biol. 2004;3:Article3.

76. Badia IMP, Velez Santiago J, Braunger J, Geiss C, Dimitrov D, Muller-Dott S, et al. decoupleR: ensemble of computational methods to infer biological activities from omics data. Bioinform Adv. 2022;2(1):vbac016.

77. Garcia-Alonso L, Holland CH, Ibrahim MM, Turei D, Saez-Rodriguez J. Benchmark and integration of resources for the estimation of human transcription factor activities. Genome Res. 2019;29(8):1363–75.

78. Holland CH, Szalai B, Saez-Rodriguez J. Transfer of regulatory knowledge from human to mouse for functional genomics analysis. Biochim Biophys Acta Gene Regul Mech. 2020;1863(6):194431.

79. Korotkevich GS, Vladimir; Budin, Nikolay; Shpak, Boris; Artyomov, Maxim N.; Sergushichev, Alexey. Fast gene set enrichment analysis. bioRxiv. 2021;10.1101/060012.

