## Supplemental Figures for "SIRT1 remodels astrocyte metabolism and promotes viral replication during neurotropic orthoflavivirus infection"

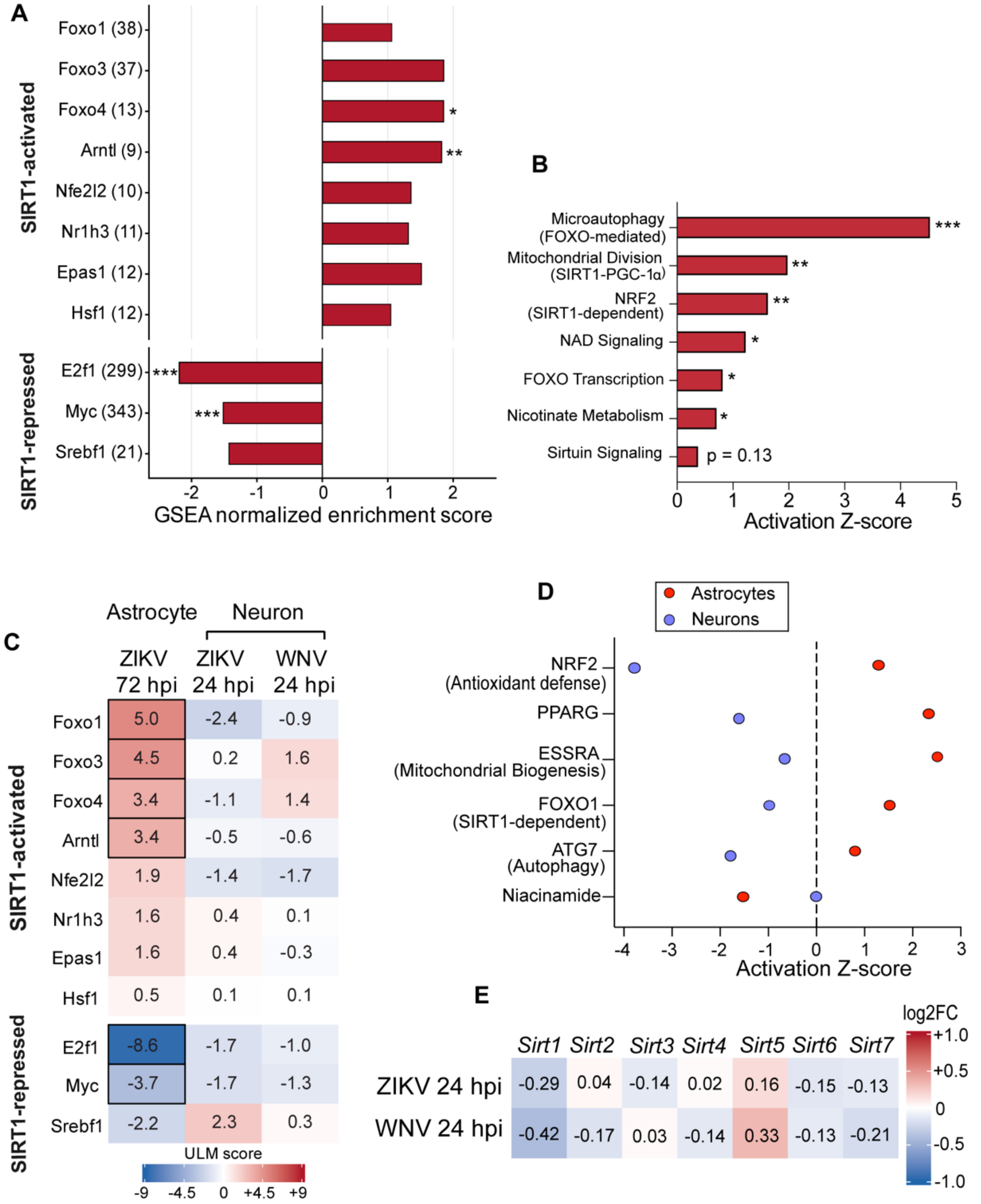


**Supplemental Figure 1. Flavivirus infection engages a SIRT1-associated transcriptional program in astrocytes but not in neurons.** (A-B) Gene set enrichment analysis for genes associated with the indicated regulons (A) and Ingenuity Pathway Analysis showing canonical pathways enriched (B) in ZIKV-infected primary mouse astrocytes at 72 hpi relative to mock. (C) Transcription factor activity for the SIRT1-substrate regulons in primary mouse cortical neurons infected with ZIKV or WNV at 24 hpi, inferred by decoupleR from mouse DoRothEA regulons. (D) IPA Upstream Regulator predictions for the SIRT1-regulated factors NRF2, PPARG, ESRRA, FOXO1, and ATG7 in infected astrocytes versus infected neurons. (E) $Log_2$ fold change in Sirtuin-encoding gene expression in primary mouse cortical neurons infected with ZIKV or WNV at 24 hpi relative to mock. Cells outlined in black indicate padj < 0.05. Comparisons of regulon and expression analyses via Benjamini-Hochberg procedure. **p* < 0.05, ***p* < 0.01, ****p* < 0.001.

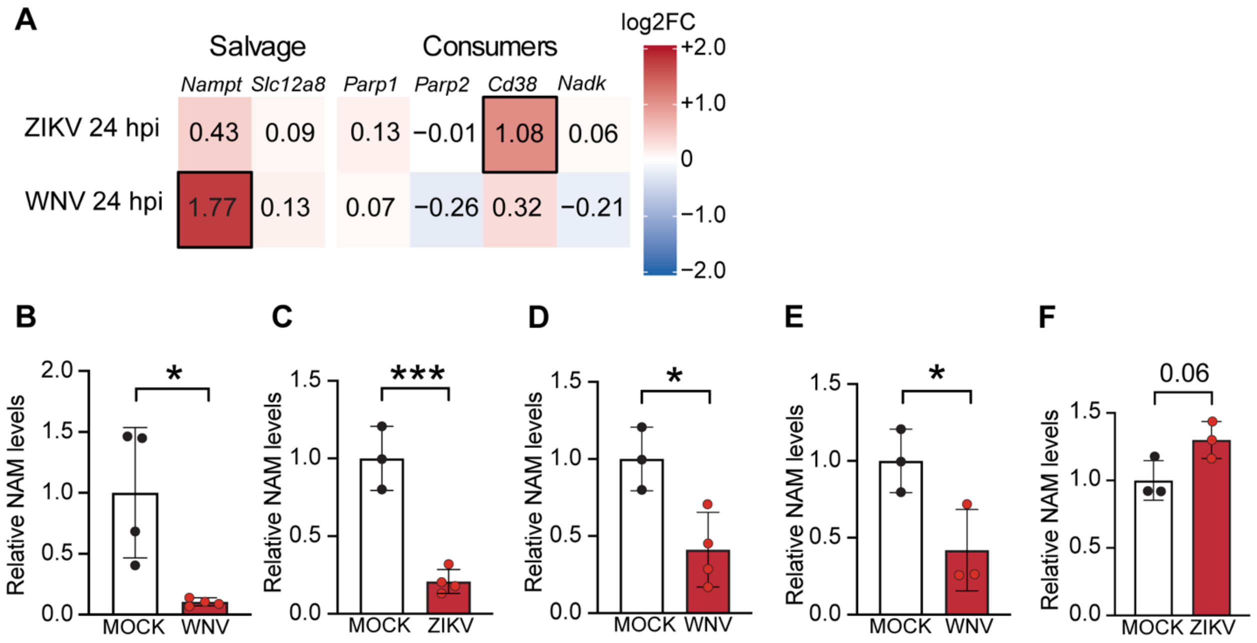


**Supplemental Figure 2. Flavivirus infection depletes nicotinamide and engages the coordinated NAD salvage response in astrocytes but not in neurons**. (A) $Log_2$ fold change in NAD+ metabolic-network associated gene expression in primary mouse cortical neurons infected with ZIKV or WNV at 24 hpi relative to mock. Cells outlined in black indicate padj < 0.05. (B) NAM abundance in whole-brain homogenates from mock- and WNV-infected mice at 9 dpi, measured by ELISA. n = 5 mice per group. (C–E) NAM abundance in cultured primary mouse astrocytes, mock-infected or virus-infected, measured by ELISA: ZIKV at 48 hpi (C), WNV at 48 hpi (D), and WNV at 72 hpi (E). n = 3 independent cultures/condition (each comparison shares the same control as infections were performed simultaneously). (F) NAM abundance in primary mouse cortical neurons, mock-infected or ZIKV-infected at 24 hpi, measured by ELISA. n = 3 independent cultures/condition. For (B–F), NAM concentrations were normalized to total protein by BCA assay. (B-F) represent mean values ± SEM. Comparisons in (B-F) via Welch's two-tailed t-test. $*p < 0.05$, $**p < 0.001$, $***p < 0.001$.

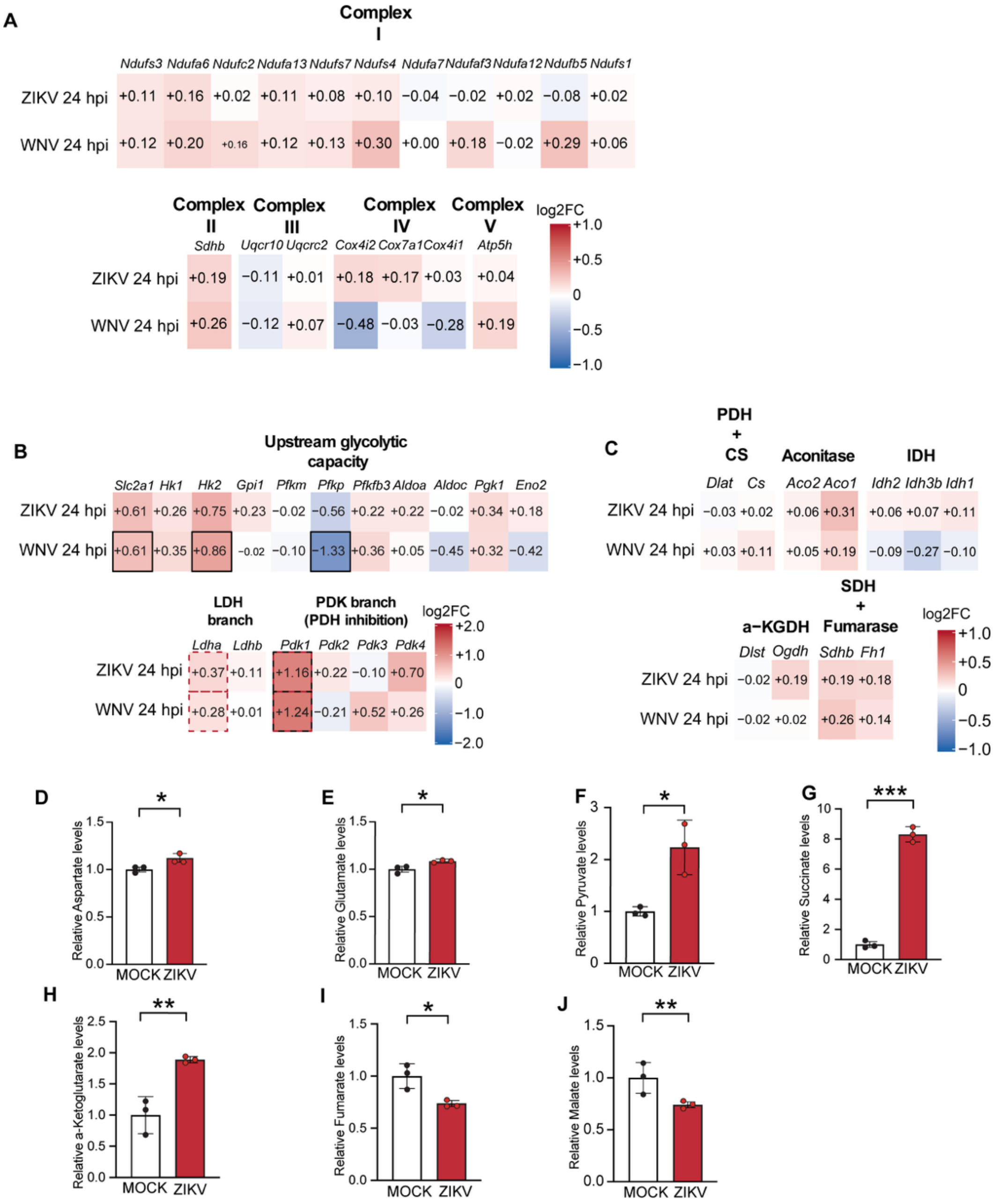


**Supplemental Figure 3. Orthoflavivirus infection in neurons produces glycolytic-associated metabolic signatures distinct from the oxidative program of astrocytes.** (A) $Log_2$ fold change in oxidative phosphorylation-associated gene expression across all five respiratory-chain complexes in primary mouse cortical neurons infected with ZIKV or WNV at 24 hpi relative to mock. Cells outlined in black indicate padj < 0.05. (B) $Log_2$ fold change in glycolysis-associated gene expression in the same neuronal dataset. Cells outlined in black indicate padj < 0.05. (C) $Log_2$ fold change in TCA cycle-associated gene expression in the same neuronal dataset. Cells outlined in black indicate padj < 0.05. (D-E) Relative levels of aspartate (D) and glutamate (E) in primary mouse astrocytes, mock-infected or ZIKV-infected at 48 hpi, measured by LC-MS. n = 3 independent cultures/condition. (F–J) Relative metabolite abundance in primary mouse cortical neurons, mock-infected or ZIKV-infected at 24 hpi, measured by LC-MS: pyruvate (F), succinate (G), α-ketoglutarate (H), fumarate (I), and malate (J). n = 3 independent cultures/condition. (D-J) represent mean values ± SEM. Comparisons in (D=J) via Welch's two-tailed t-test. *$p$ < 0.05, **$p$ < 0.001, ***$p$ < 0.001.
